# A simple protein-based surrogate neutralization assay for SARS-CoV-2

**DOI:** 10.1101/2020.07.10.197913

**Authors:** Kento T. Abe, Zhijie Li, Reuben Samson, Payman Samavarchi-Tehrani, Emelissa J. Valcourt, Heidi Wood, Patrick Budylowski, Alan P. Dupuis, Roxie C. Girardin, Bhavisha Rathod, Jenny H. Wang, Miriam Barrios-Rodiles, Karen Colwill, Allison J McGeer, Samira Mubareka, Jennifer L. Gommerman, Yves Durocher, Mario Ostrowski, Kathleen A. McDonough, Michael A. Drebot, Steven J. Drews, James M. Rini, Anne-Claude Gingras

## Abstract

Most of the patients infected with severe acute respiratory syndrome coronavirus 2 (SARS-CoV-2) mount a humoral immune response to the virus within a few weeks of infection, but the duration of this response and how it correlates with clinical outcomes has not been completely characterized. Of particular importance is the identification of immune correlates of infection that would support public health decision-making on treatment approaches, vaccination strategies, and convalescent plasma therapy. While ELISA-based assays to detect and quantitate antibodies to SARS-CoV-2 in patient samples have been developed, the detection of neutralizing antibodies typically requires more demanding cell-based viral assays. Here, we present a safe and efficient protein-based assay for the detection of serum and plasma antibodies that block the interaction of the SARS-CoV-2 spike protein receptor binding domain (RBD) with its receptor, angiotensin converting-enzyme 2 (ACE2). The assay serves as a surrogate neutralization assay and is performed on the same platform and in parallel with an enzyme-linked immunosorbent assay (ELISA) for the detection of antibodies against the RBD, enabling a direct comparison. The results obtained with our assay correlate with those of two viral based assays, a plaque reduction neutralization test (PRNT) that uses live SARS-CoV-2 virus, and a spike pseudotyped viral-vector-based assay.

## Introduction

The coronavirus S-protein (spike) is responsible for both receptor binding and fusion of the virus and host cell membranes. Within the spike protein, the receptor binding domain (RBD) mediates the interaction with the host cell receptor and sequence/structural variation in the RBD is responsible for the receptor binding specificity shown by those coronaviruses that use host proteins as receptors (1).

SARS-CoV-2, like SARS-CoV, uses the cell surface carboxypeptidase angiotensin-converting enzyme 2 (ACE2) as a receptor for viral entry (**Figure 1A**). The use of a common receptor is consistent with the fact that the two viruses share a high degree of sequence similarity and that their RBDs are ~74% identical, though the RBD of SARS-CoV-2 binds ACE2 with higher affinity than does that of SARS-CoV (2). The spike proteins of both viruses are also both primed by the host protease, TMPRSS2, but unlike SARS-CoV-2, the spike protein of SARS-CoV does not contain a furin recognition motif that can be cleaved during viral biogenesis (2, 3).

**Figure 1.**
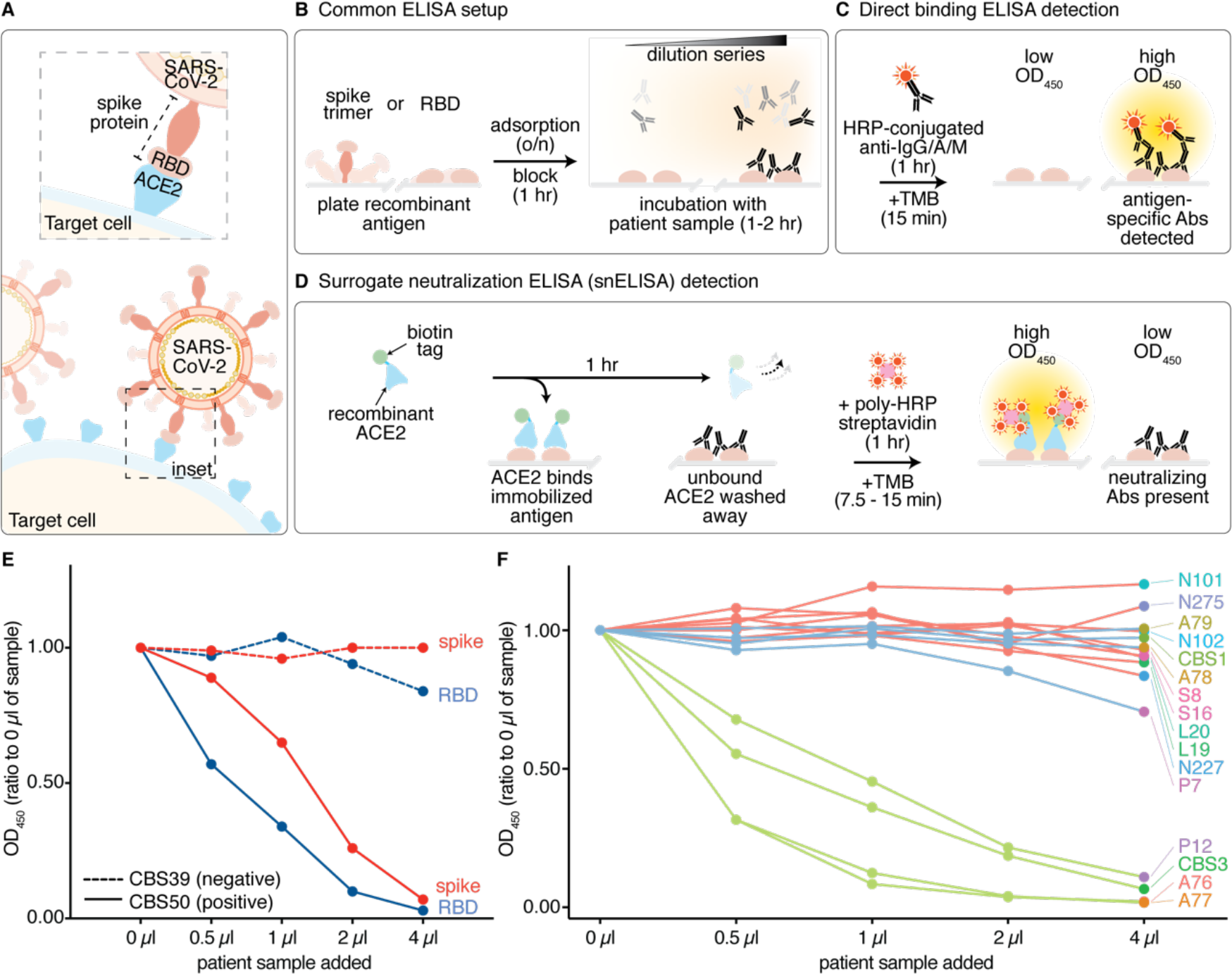
Establishment of a surrogate neutralization ELISA (snELISA) to monitor the spike-ACE2 interaction. (**A**) SARS-CoV-2 attachment to the host cell requires a direct interaction between the host cell receptor, ACE2 (blue), and the Receptor binding domain (RBD) of the SARS-CoV-2 spike protein (peach). (**B**) 96-well plates are set up in a similar manner for both the detection of anti-SARS-CoV-2 antibodies and neutralizing antibodies. Antigens are adsorbed overnight and incubated with diluted patient serum/plasma samples, monoclonal antibodies (Ab) and other affinity reagents. Antigenspecific antibodies are coloured black. (**C**) Principle of direct binding ELISA detection using HRP-conjugated anti-human IgG/A/M. (**D**) Principle of the snELISA, which uses biotinylated ACE2 for the detection of RBD or spike epitopes that have not been blocked by neutralizing antibodies (using polyHRP-streptavidin). (**E**) Results of the snELISA assay using either the RBD or the spike trimer immobilized on the plate (See **Supplemental Figure 1-2** for the antigen cloning, expression and purification). The dashed lines are from a sample (CBS39) that was negative for direct RBD/spike binding, while the solid lines are from a positive sample (CBS50; See **Supplemental Figure 4A** for the direct binding results and **Supplemental Table 1** for all OD_450nm_ values). (**F**) snELISA (immobilized RBD) for an expanded set of 4 positive controls with high anti-RBD signals in a single-point direct-binding ELISA (green), 8 negative samples acquired pre-COVID (red) and 4 samples with low anti-RBD levels (blue; **Supplemental Figure 4A**). The five-point curves in **E-F** were generated from one experiment.

The coronavirus spike protein is also a major target of the host immune system and antibodies directed against it play a central role in host-mediated neutralization (4). Among neutralizing antibodies, those that block the interaction between viruses and their receptors represent the most common route to neutralization (5). For this reason, both the spike protein and the RBD form the basis for most of the SARS-CoV-2 vaccines currently in development. The detection and study of neutralizing antibody activity following natural infection (or vaccination) can, therefore, support research aimed at the development of novel therapeutics and vaccine candidates. It can also aid in the identification of acceptable donors for convalescent plasma therapy (6), and more generally to establish immune correlates of infection.

For SARS-CoV-2, viral neutralization assays are performed using either live virus [e.g. (7)] or viral vectors pseudotyped with the spike protein [e.g. (8)]. However, these cell culture-based assays are challenging to implement and time-consuming to run, factors that limit scalability. The conventional plaque reduction neutralization test (PRNT) that uses live SARS-CoV-2 virus is further complicated by the need for containment level 3 (CL-3) and a specialized laboratory setup. Although the pseudotyped viral-vector-based assays do not require biosafety level 3 (BSL-3) containment (8), they are nevertheless complicated multistep procedures (9).

By contrast, the detection and quantitation of antigen-specific antibodies in patient samples can be easily assayed by enzyme-linked immunosorbent assays [ELISA, see e.g. (10)]. SARS-CoV-2 ELISAs are performed by immobilizing a recombinantly produced viral antigen (such as the spike trimer or RBD; **Figure 1B, Supplemental Figures 1, 2**; see Methods) onto multi-well plastic plates that are then incubated with diluted patient serum or plasma samples. The detection of antibodies that bind to the antigen involves a second incubation with enzyme-conjugated anti-human antibodies, where the enzyme is often horseradish peroxidase (HRP). This enables the detection of a color change when an HRP substrate such as 3,3’,5,5’-tetramethylbenzidine (TMB) is used. In direct binding assays of this type (**Figure 1C**), the presence of patient antibodies against the viral antigen leads to a dosedependent increase in the signal observed.

ELISA-based profiling has been developed by multiple groups and used to measure the kinetics of the antibody response in patient cohorts following SARS-CoV-2 infection. In several recent studies, including ours, this has revealed the relative stability in the IgG response to the spike and RBD over several months, and a more transient IgM and IgA response that wanes as patients convalesce (11-15). However, the levels of *neutralizing* antibodies are not typically measured in large cohorts over time [with a few notable exceptions, e.g. (12, 15)], as current assays are relatively low throughput. The relative lack of neutralizing antibody data represents a significant gap in our understanding of the immune response to SARS-CoV-2.

Here, we describe a modified ELISA-type assay that serves as a surrogate neutralization assay. It measures the presence of antibodies capable of blocking the RBD-ACE2 interaction and, like the direct binding ELISA, it is easily scaled to allow for the analysis of large patient cohorts over time. We show that the results obtained by this assay correlate with those of both the SARS-CoV-2 PRNT and a spike pseudotyped viral-vector neutralization assay in a cohort of convalescent patients and on purified antibodies.

## Results

We aimed to develop a simple protein-based assay to monitor the ability of antibodies, present in the serum or plasma of patients, to block the interaction between the RBD and the host receptor ACE2. To do so, we elected for an ELISA-type assay since such assays are already widely used to detect antibodies that recognize SARS-CoV-2 antigens such as the spike trimer and its RBD. As with the standard direct ELISA, the antigen (here, the RBD or the spike trimer) is first immobilized on multi-well plates and then incubated with patient plasma or serum **(Figure 1B)**. However, because we were interested in detecting functional antibodies that can prevent the interaction between the RBD (or spike) and ACE2, we replaced the HRP-conjugated secondary antibody used in the direct ELISA by a detection method involving human ACE2. In our assay, recombinantly-expressed soluble ACE2 bearing a biotinylated C-terminal AviTag is added to the antigen-bound plate after the plate has been incubated with the patient plasma or serum (see Methods). Bound ACE2 is then detected by the addition of streptavidin-poly HRP and its colorimetric substrate TMB. The presence of patient antibodies that can block the RBD-ACE2 interaction leads to a dose-dependent decrease in the signal observed and, as such, we refer to it as a surrogate neutralization ELISA assay (*snELISA* - **Figure 1D**).

We explored two different versions of the assay: the configuration described above and one involving immobilized ACE2 and soluble biotinylated RBD, a configuration similar to that previously reported (16). In our hands, the assay with immobilized RBD and soluble biotinylated ACE2 was more sensitive than its counterpart (**Supplemental Figure 3**). Moreover, with the RBD immobilized, the same overall protocol and colorimetric detection can be used for both the direct binding ELISA and the snELISA, thereby facilitating a direct comparison.

Although the snELISA worked well with either the RBD or the spike ectodomain trimer immobilized (**Figure 1E**; **Supplemental Figure 4A**), we focused on the RBD as it is easier to produce and provides a simple one-to-one binding interaction with ACE2. Using a small test set (**Supplemental Figure 4B**), we first showed that the serum/plasma from positive, but not negative control patients, inhibited the interaction between ACE2 and the immobilized RBD (**Figure 1F**). The technical reproducibility of the assay was within 5–10 % CV. The total time required to perform the assay (once the plates are coated with the antigen) is 3.5–4 hours, and the assay can be performed using the same equipment and biosafety protocols as a standard ELISA.

Using both the surrogate neutralization and direct binding (with a dilution series) ELISA assays, we then profiled a set of 58 serum samples acquired at the Canadian Blood Services as part of a screen for convalescent plasma therapy donors (**Figure 2A–B; Supplemental Figures 5–8**; **Supplemental Table 1**). With reference to the direct binding results, the snELISA showed that samples with high levels of IgG against the RBD were typically the most potent at blocking the RBD-ACE2 interaction (e.g. CBS13, which is included as a positive control). Conversely, samples lacking detectable RBD-binding antibodies were not able to block the interaction.

**Figure 2.**
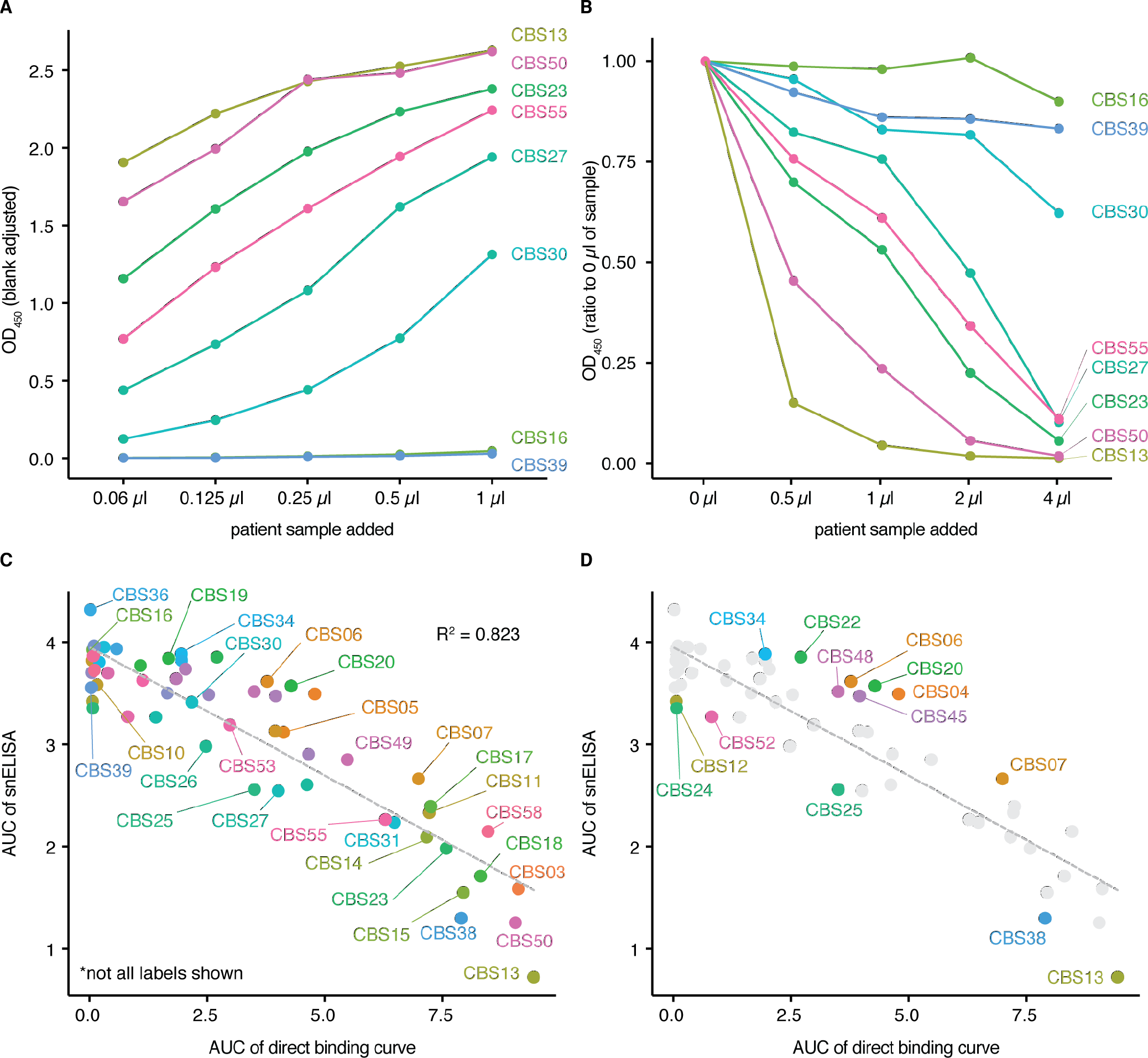
Application of the snELISA to a larger cohort. (**A**) Representative direct binding ELISAs with titrations of different samples from a patient cohort sampled by Canadian Blood Services (all 58 ELISA curves are shown in **Supplemental Figure 6**; see **Supplemental Figure 7** for an extended titration of the most abundant samples). (**B**) snELISA results for the samples shown in **A**, see **Supplemental Figure 8** for all curves. The five-point curves in **A-B** were generated from one experiment. (**C**) Correlation between the Areas Under the Curves (AUCs) for the direct and snELISAs for all samples profiled (see an expanded view in **Supplemental Figure 9**). (**D**) Outliers in the correlation plot **C** were calculated using the total least squares (TLS) method; points with a TLS > 0.4 (labelled) were marked as outliers (See **Supplemental Figure 10** for selected examples with side-by-side direct and snELISAs).

To more systematically evaluate the relationship between the RBD-binding antibody levels and the ability to block the RBD-ACE2 interaction (as determined by the snELISA), we calculated the area under the curve (AUC) for both assays and plotted the RBD-binding AUC versus the snELISA AUC (**Figure 2C; Supplemental Figure 9**). The plot showed a clear correlation (R^2^ = 0.823), with the sera containing the highest RBD-binding antibody levels being the most effective at blocking the RBD-ACE2 interaction (**Figure 2C**; also compare **Figure 2B** to **2A**; **Supplemental Table 1**). Nevertheless, there are samples with similar RBD-binding antibody concentrations that differ in their ability to block the RBD-ACE2 interaction (**Figure 2D; Supplemental Figure 10**). Differences in antibody isotype, affinities, and abundance, as well as the RBD epitopes bound, are all factors that could explain these outliers.

While it is reasonable to expect that antibodies that block the RBD-ACE2 interaction would be neutralizing, we validated this using cell-based viral infectivity and entry assays. Fifty-seven of the 58 samples analyzed by the snELISA were analyzed by PRNT, the gold standard in the field. PRNT50 is defined as the concentration of patient serum or plasma capable of reducing the formation of viral plaques by 50% (PRNT90 is the concentration that reduces plaque formation by 90%). As shown in **Figure 3A,** most of the samples displaying high values in the direct binding and snELISA assays were also positive by PRNT90 (and those with low titers were negatives). Both ELISA assays also gave an overall agreement with the PRNT50 titers (see **Supplemental Figure 11**, with a coefficient of determination of 0.6). We also adapted and optimized a spike-pseudotyped lentiviral-based entry assay (8), and reprofiled the neutralization potential of a subset of samples. There was also a high correlation (R^2^ = 0.76) between the snELISA assay results and the titers obtained with this spike-pseudotyped lentiviral-based entry assay (**Figure 3B** and **Supplemental Figure 12**). Taken together, these results indicate that our snELISA is a good surrogate neutralization assay, particularly for distinguishing between samples with high versus low neutralization activity. As such, the assay should be of value in the selection of candidate donors for convalescent plasma therapy and for monitoring immune correlates of patient outcomes. Future work will focus on providing a better understanding of the outliers observed across all assays. Indeed, rare but potent neutralizing antibodies in patient samples with low pseudovirus neutralization titers have recently been reported (17).

**Figure 3.**
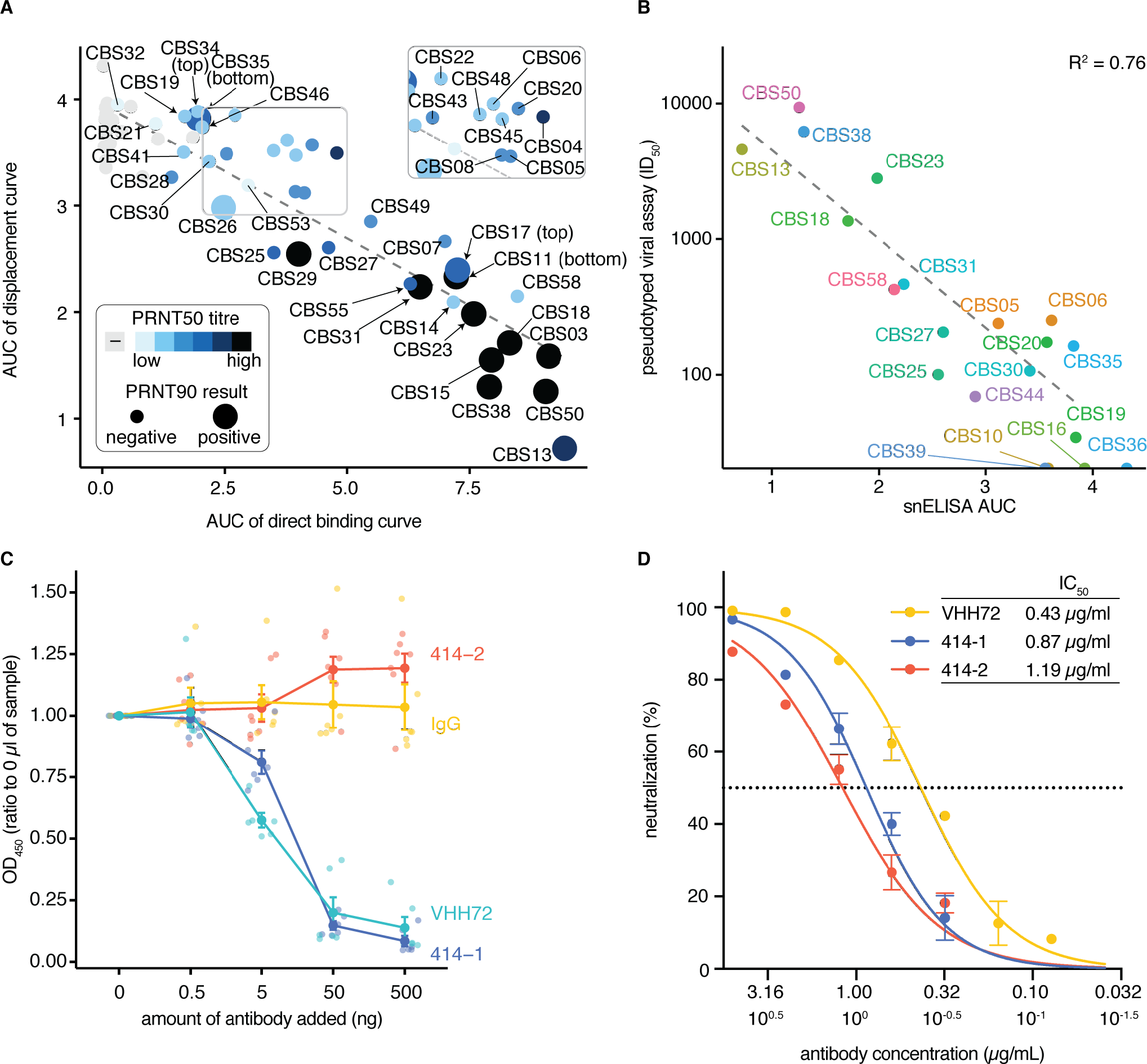
Validation of the snELISA results using orthogonal methods. (**A**) Results of the plaque reduction neutralization tests on the same samples overlaid on the AUC curves from **Figure 2C** (n=57 samples). Color coding indicates the PRNT50 titers while negative/positive hits on the PRNT90 assay are displayed with a different sized dot (see **Supplemental Figure 11** for additional PRNT results and **Supplemental Figure 12** for spike pseudotyped virus results). (**B**) Correlation between the lentiviral spike pseudotyped virus assay calculated IC_50_ values and the AUC results from the snELISA. Assessment of the ability of monoclonal or affinity reagents to block the interaction between ACE2 and the RBD in (**C**) the snELISA or (**D**) the lentiviral spike pseudotyped assay (see **Supplemental Figure 13-16** for the direct binding and viral neutralization assays and additional reagents tested). Values for panel **C** were generated from three independent experiments, and underlayed points have been jittered along the x-axis. Values for panel **D** were obtained in one experiment in technical triplicate. Error bars represent the mean ± standard error of the mean (SEM).

To assess whether our snELISA might also be of value for screening the neutralization potential of monoclonal antibodies, we tested it using a number of neutralizing and non-neutralizing monoclonal antibodies and compared the results with that obtained with the pseudotyped lentiviral-based entry assay or cytopathic effect-reduction neutralization assay with SARS-CoV-2. The llama VHH72 monoclonal antibody (expressed as a human Fc fusion), previously shown to neutralize in a SARS-CoV-2 spike pseudotyped entry assay (18), blocked the RBD-ACE2 interaction in our snELISA and viral entry in our spike pseudotyped lentivirus assay; similar results were obtained for the Active Motif 414-1 antibody which was isolated from a convalescent patient and shown to be neutralizing (19) (**Figure 3C, 3D; Supplemental Figure 13**). In contrast, other antibodies such as an IgG derived from the monoclonal anti-SARS1 CR3022 or a commercial antibody from GenScript, had a much more moderate effect in the snELISA (**Supplemental Figure 14**) and the Genscript antibody had no effect in the cytopathic effect-reduction neutralization assay (**Supplemental Figure 15**). The Active Motif 414-2 antibody was previously shown to be incapable of neutralizing live SARS-CoV-2 virus (8, 19). In our assays, it efficiently bound the RBD in the direct ELISA but did not block the RBD-ACE2 interaction in the snELISA. The same antibody partially prevent entry in our lentivirus entry assay. Taken together, these observations suggest that our surrogate neutralization ELISA is a good complement to more complex cell-based assays for the discovery and screening of neutralizing monoclonal antibodies.

## Discussion

In summary, we have developed a simple and safe surrogate neutralization ELISA for SARS-CoV-2. It can be readily incorporated into existing testing platforms and may be of particular value in the selection of donors for convalescent plasma therapy and as a means of monitoring the immune response to vaccination. Given that neutralizing antibody titres have recently been shown to wane fairly rapidly in some (12-14, 20) but not all (15, 20) studies, the assay may also be useful for broad serosurveillance, especially as it should be more scalable than the approaches requiring viral infection assays. When coupled with epidemiological studies, it might also be used to assess the risk of infection/re-infection. We also note that the optimized conditions used here for the direct RBD-binding ELISA are similar to those reported by the Krammer lab using RBD-expression constructs that have been widely distributed. Their RBD can be obtained from BEI resources (10), and we found that it generates similar results when used with our biotinylated ACE2 in the snELISA (**Supplemental Figure 16**). This should further facilitate the broad implementation of our assay across multiple laboratories.

There are limitations to the assay, however, that need to be acknowledged. First, the snELISA is limited to the detection of neutralizing antibodies that function by blocking the interaction between the RBD and ACE2. While by no means dominant, examples of antibodies that neutralize by other mechanisms are beginning to emerge (21-23). The snELISA, in conjunction with a neutralization assay, could be used to identify further such examples. As with those identified in this work, the outliers (e.g. those with high viral neutralization titers but low snELISA levels) provide a starting point for further work aimed at understanding the mechanisms of antibody-mediated neutralization.

Another limitation of our approach is that the current assay cannot directly map the epitopes targeted by the various antibodies. Undoubtedly, the antibodies detected by the snELISA bind to different sites on the RBD, a suggestion supported by the structures of neutralizing antibody Fabs in complex with the SARS-CoV-2 RBD. In one example, two different neutralizing antibodies that bind to different epitopes on the RBD were found to synergistically mediate viral neutralization (24). While in the current study we simply wanted to provide evidence of antibodies that could block the RBD-ACE2 interaction, the snELISA could be adapted to provide information on the site of antibody binding. As recently shown, a series of structure-guided point mutants in the RBD could be used to infer where on the RBD the antibodies are binding (25). This type of approach would likely be more important in the characterization of monoclonal antibodies, such as those presented in **Figure 3C**, and would set the stage for in depth biophysical and structural studies.

While the direct binding ELISA described here employed an anti-IgG secondary antibody (the predominant isotype in convalescent serum), we note that the snELISA measures the ability of any antibody isotype (or even antibodies from different species or any other molecule) to block the RDB-ACE2 interaction. In this regard, it is similar to that of a viral-based neutralization assay. While we have not performed a detailed analysis, we did show that single point direct binding ELISAs performed for IgM, and to a lesser extent IgA, are also correlated with the results obtained by the snELISA (**Supplemental Figure 17**). The safety and simplicity of the snELISA should make it a valuable addition to the arsenal of assays for monitoring the immune response to SARS-CoV-2 infection.

## Methods

### Serum and plasma samples

#### Canadian Blood Services Donors

Specimens-only serum donations were collected from individuals who were classified as having one or more of the following criteria: 1) self-declared evidence of a SARS-CoV-2-positive nucleic acid test, 2) a declaration of having been a close contact of a COVID-19 case, 3) a travel history and clinical presentation compatible with COVID-19, and 4) signs and symptoms compatible with COVID-19. Collections occurred two weeks or more after cessation of clinical symptoms. Serum specimens were processed and frozen at −80°C until shipment on dry ice to the testing laboratory.

#### Other samples for assay development

Negative control serum samples from patients enrolled in cancer studies prior to COVID-19 (prior to November 2019; REB studies #01-0138-U and #01-0347-U, Mount Sinai Hospital) and archived frozen in the LTRI Biobank were retrieved, thawed, aliquoted and transferred on ice to the research lab for viral inactivation. Alternatively, samples from previous studies of the immune system or systemic lupus acquired prior to November 2019 (REB studies #31593 University of Toronto, #05-0869, University Health Network, a kind gift from Dr. Joan Wither, University Health Network, Toronto, ON, Canada) were transferred to the lab on dry ice. Positive controls for assay development were either convalescent plasma or serum from COVID-19 patients (confirmed by PCR; in- and out-patients) acquired in south-central Ontario in 2020 (REB studies #20-044 Unity Health Network, #02-0118-U/05-0016-C, Mount Sinai Hospital). Aliquots of these samples were transferred to the lab on dry ice. Only those samples with high levels of RBD-binding antibodies in single-point ELISA assays were considered positives for the development of the snELISA. Samples with low levels of RBD-binding antibodies were reclassified as “negative”.

For all ELISAs, inactivation of potential infectious viruses in plasma or serum was performed by incubation with Triton X-100 to a final concentration of 1% for 1 h prior to use (26). For the pseudotyped lentiviral assays, the serum was heat-inactivated for 1 hr at 56° C (10). See the different versions of the PRNT assays for details of the inactivation procedures, as applicable.

### Expression system for protein purification

The expression plasmid generated is a derivative of those previously reported in our piggyBac transposon-based mammalian cell expression system (27). Two versions of the plasmid were constructed; one contains the CMV promoter (PB-CMV) and the other the TRE promoter (PB-TRE). The vectors are otherwise identical and can be used to generate stable cell lines for constitutive or inducible protein expression. The protein cloning region contains several optional elements separated by restriction sites as follows: an N-terminal human cystatin-S secretion signal, the protein of interest, a foldon trimerization motif (28), a 6xHis purification tag and an AviTag biotinylation motif (29) (**Supplemental Figure 1**). A woodchuck hepatitis virus posttranscriptional regulatory element (WPRE) follows the ORF to facilitate nuclear export of the mRNA. A pair of piggyBac transposon terminal repeats flank the expression cassette and an attenuated puromycin resistance marker, thereby allowing for the generation of stable cell lines using the piggyBac transposase.

### Expression constructs for protein purification

The human codon optimized cDNA of the SARS-CoV-2 spike protein (MC_0101081) was purchased from GenScript (Piscataway, NJ, USA). The human ACE2 cDNA was derived from MGC clone 47598. To stabilize the soluble spike ectodomain trimer, two regions of the spike protein were mutated. Residues 682–685 (RRAR) were mutated to SSAS to remove the furin cleavage site, and residues 986– 987 (KV) were each mutated to a proline residue to stabilize the pre-fusion form as previously described (30). The soluble spike protein ectodomain construct includes residues 1-1211 (MFVF…QYIK), followed by the foldon trimerization motif, a 6xHis tag and an AviTag. Both the SARS-CoV-2 receptor binding domain (RBD) and the human ACE2 constructs are preceded by the human cystatin-S secretion signal and followed by the 6xHis and AviTag. The RBD and ACE2 constructs contain residues 328-528 (RFPN…CGPK) and 19-615 (STIE…PYAD), respectively.

The cDNA of the human CR3022 Fab fragment was synthesized by GenScript based on its previously reported sequence (31). The light chain and heavy chains were individually cloned into the PB-TRE expression plasmid. For Fab production, a 6xHis tag was added to the C-terminal end of the Fab heavy chain. An IgG form was generated by fusing the human IgG1 Fc coding sequence to the C-terminal end of the Fab heavy chain.

### Large scale transient transfection

FreeStyle 293-F suspension cells were grown in shaker flasks (125 rpm) in Freestyle 293 expression medium (ThermoFisher) in a humidified 37 °C incubator filled with 3% (v/v) CO2. The cell density and viability were monitored by manual counting using a hemocytometer and trypan blue staining. For transfection, cells of >90% viability were counted and seeded at a density of approximately 10^6^ cells/mL into 300 mL Freestyle 293 medium supplemented with 1 μg/mL Aprotinin. 300 μg of the PB-CMV plasmid DNA and 400 μL 293fectin (ThermoFisher) were each added to separate tubes containing 15 mL of Opti-MEM medium (ThermoFisher). The two solutions were then mixed and incubated for 5 min before being added to the 300 mL cell culture. Two days after transfection, the 300 mL culture was expanded into three 1L shaker flasks each containing 300 mL of culture medium. Protein expression was continued for an additional four days.

### Stable cell line generation

FreeStyle 293-F cells or a GnT1-knockout FreeStyle 293-F cell line were used for generating stable cell lines. Approximately 10^6^ cells were added to each well of a 6-well plate in 2 mL FreeStyle 293 medium. 2 μg of the PB-TRE plasmid encoding the protein of interest, 0.5 μg of the PB-rtTA-neomycin helper plasmid(27) and 0.5 μg of the PBase expression plasmid, pCyL43(32), were co-transfected in each well using lipofectamine 2000 (Thermofisher) following the manufacturer’s instructions. Three days post-transfection, the cells were transferred to 10-cm dishes containing FreeStyle 293 medium supplemented with 10% FBS, 2 μg/mL puromycin and 200 μg/mL G418. Selection was continued for approximately 2 weeks.

The stable cells were scaled up in 1L shaker flasks containing 300 mL FreeStyle 293 medium without supplements. When the cell densities reached approximately 10^6^ cells/mL, 1 μg/mL doxycyline and 1 μg/mL Aprotinin were added to initiate protein expression. During the expression phase, 150 mL of the medium was removed, and fresh medium added, every other day.

### Protein purification

The harvested expression medium was centrifuged at 10,000*g* to remove the cells and debris. For the 6xHis tagged proteins, the clarified media were passed through an Ni-NTA column. For the spike ectodomain, 3 mL of Ni-NTA resin was used for each liter of medium. For the RBD, ACE2 and CR3022 Fab, 8 mL of Ni-NTA resin was used for each liter of medium. The Ni-NTA resin was washed with 20 column volumes of phosphate buffered saline (PBS), followed by 3–5 column volumes of PBS containing 10 mM imidazole. The protein was eluted with PBS containing 300 mM imidazole and 0.1% (v/v) protease inhibitor cocktail (Sigma, P-8849). For the CR3022 antibody, the harvested medium was incubated with rProtein A Sepharose FF resin (GE healthcare). The resin was then washed with 20 column volumes of PBS and the antibody was eluted with 50 mM glycine, pH 3.0, containing 150 mM NaCl. The acid-eluted antibody was immediately neutralized by the addition of 1/20 volume of 1 M Tris, pH 8.5. Protease inhibitor cocktail was also added to a final concentration of 0.1% (v/v). The approximate purified yields of the various proteins are as follows: RBD, 70 mg/L; spike trimer, 3 mg/L; ACE2, 50 mg/L; CR3022 Fab, 80 mg/L and CR3022 IgG, 20 mg/L.

The protein samples were stored in 40% glycerol at −12 °C. Shortly before use, the glycerol stocks were further purified using size-exclusion chromatography. For the RBD, ACE2 and CR3022 Fab/IgG, a Superdex 200 Increase (GE healthcare) column was used. For the spike ectodomain, a Superose 6 Increase (GE healthcare) column was used (**Supplemental Figure 2**).

### Site-specific biotinylation of the AviTag containing proteins

Each biotinylation reaction contained 200 μM biotin, 500 μM ATP, 500 μM MgCl2, 30 μg/mL BirA, 0.1% (v/v) protease inhibitor cocktail and no more than 100 μM of the protein-AviTag substrate. The mixture was incubated at 30 °C for 2 hr followed by size-exclusion chromatography to remove unreacted biotin. For the RBD, the degree of biotinylation was assessed using a band-shift assay. 5 μg of the biotinylated RBD was heated to 95 °C for 30 s in SDS-PAGE loading buffer (containing 2% SDS, 50 mM DTT) and then after cooling, 1 μL of a 5 mg/mL streptavidin solution was added. The mixture was then analyzed by SDS-PAGE to assess the formation of the RBD-streptavidin complex (**Supplemental Figure 2**).

### Production of the VHH72 recombinant antibody

The llama single domain antibody VHH72 sequence (PDB entry 6WAQ_1) was obtained from Wrapp *et al.* (18). A cDNA encoding VHH72 fused to an ADCC-attenuated human IgG1 Fc domain (hFc1X7, from patent US 2019 352 383A1) was codon-optimized for expression in *Cricetulus griseus* (CHO cells), synthesized by GenScript and cloned into the pTT5™ plasmid (33). The pTT5-VHH72hFc1X7 plasmid was transiently expressed in CHO^55E1^ cells (34) using PEI-Max transfection reagent (Polysciences, Warrington, PA) and a slightly modified protocol as described previously (35). The cell culture was harvested at day 7 post-transfection, centrifuged 20 min at 3000*g* and filter-sterilized using a 0.22 μm membrane vacuum filter (Express PLUS, Millipore). Filtered supernatant was loaded on a 5 ml MabSelect SuRe column (GE Healthcare) equilibrated in PBS. The column was washed with PBS and the antibody eluted with 100 mM citrate buffer pH 3.6. The fractions containing the antibody were pooled and elution buffer was exchanged for PBS using NAP-25 columns (GE Healthcare). Purified VHH72hFc1X7 in PBS was quantified by absorbance at 280 nm using a Nanodrop spectrophotometer (Thermo Fisher Scientific) and the calculated extinction coefficient of the protein. Overall volumetric yield post-protein-A purification was 275 mg/L. The purified protein was analyzed by analytical size-exclusion ultra-high performance liquid chromatography coupled to a MALS detector and eluted as a major (>98% integrated area) symmetrical peak of 102 kDa with less than 2% aggregates (not shown).

### Sources of other commercial proteins and recombinant antibodies

An alternative source for RBD was BEI Resources NR-52306 (contributors F Krammer, F Amanat, S Strohmeier; lot #7034437). Commercial antibodies tested also included a human IgG chimeric antibody from GenScript (SARS-CoV-2 spike S1 Antibody (HC2001), GenScript #A02038) and two SARS-CoV-2 spike Antibodies from Active Motif (AM002414, #91349; AM001414, #91361).

### Direct ELISA assay for the identification of antibodies to the RBD

*<u>For the manual single point ELISAs in 96-well format</u>,* concentrations and incubation times were optimized to maximize the separation between anti-RBD levels in convalescent plasma or serum from that of pre-COVID era banked serum while maintaining the required levels of antigens as low as possible. One microliter of serum or plasma was used for the detection of antibodies on 96-well plates coated with 75 ng/well of recombinant purified RBD. Single point ELISAs are expressed as ratios to a positive control convalescent plasma sample. *<u>For the multipoint ELISAs</u>,* the RBD amount was fixed to 100 ng/well to match the design of the snELISA, and two-fold serial dilutions of the serum or plasma sample from 1 μl to 0.06 μl were employed.

*<u>In both cases</u>,* the RBD antigen (diluted to 2 μg/ml in PBS) was first adsorbed to 96-well clear Immulon 4 HBX (Thermo Scientific, #3855) plates in PBS overnight at 4 °C, then washed three times with 200 μl PBS+ 0.1% Tween-20 (PBS-T; Sigma). Plates were blocked for 1 hr at room temperature with 200 μl 5% Blocker™ BLOTTO (Thermo Scientific, #37530) and washed three times with 200 μl PBS-T. In the single point ELISAs, plate blocking was performed with 3% w/v milk powder (BioShop Canada Inc., #ALB005.250, lot #9H61718) in PBS for 1–2 hr.

Patient samples (pre-treated with 1% final Triton X-100 for viral inactivation) diluted in PBS-T containing 1% w/v milk powder (1:50 for the single point ELISA) were then added to the plates and incubated for 2 hr at room temperature (50 μl total volume): technical duplicates were performed unless otherwise indicated. A chimeric human anti-spike antibody (SARS-CoV-2 spike S1 Antibody (HC2001), Genscript #A02038) was added to a set of wells on each plate as a serial dilution (1:5,000 to 1:80,000 or 10 ng to 0.63 ng per well in four steps) to enable cross-plate comparisons. Positive (convalescent plasma from a single patient) and negative controls (pre-COVID era banked serum) were also added to each plate, at 1 μl.

Wells were washed three times with 200 μl PBS-T. Goat anti-human anti-IgG (Goat anti-human IgG Fcy-HRP, Jackson Immunoresearch, #109-035-098) at a 1:60,000 dilution (0.67 ng/well) in 1% BLOTTO was added and incubated for 1 hr. Wells were washed three times with 200 μl PBS-T, and 50 μl of 1-Step™Ultra TMB-ELISA Substrate Solution (ThermoFisher, #34029) was added for 15 min at room temperature and the reaction was quenched with 50 μL stop solution containing 0.16N sulfuric acid (ThermoFisher, #N600). The plates were read in a spectrophotometer (BioTek Instruments Inc., Cytation 3) at 450 nm. For all ELISA-based assays, raw OD values had blank values subtracted prior to analysis. For the single-point direct binding assay, the average CV across CBS samples is 3.3% (mean) and 1.8% (median) (**Supplemental Table 1**). For single point assays, all data were normalized to the positive serum control (single point) on each plate and expressed as a ratio to this control. For the multi-point dose responses, blank-adjusted reads were used.

#### Variations to this protocol included

1. Replacement of the RBD on the plate by the BEI Resources #NR-52306. The assay was set up identically to and in parallel with our in-house produced RBD (**Supplemental Figure 16**).
2. Replacement of RBD (100 ng) on plate by the spike trimer purified above (667 ng) (**Supplemental Figure 3**).
3. Performing the single point ELISA assays using an automated platform with chemiluminescent detection for anti IgG, IgA and IgM, exactly as described in (11) (**Supplemental Figure 17**).

### Surrogate neutralization ELISA assay for the identification of neutralizing antibodies

Our final optimized snELISA assay used 100 ng immobilized recombinant RBD on 96-well Immulon HBX plates incubated overnight at 4 °C (2 μg/ml). All volumes added to the well were 50 μl, unless specified otherwise. Plates were washed three times with 200 μl PBS-T and blocked for 1-1.5 hr at room temperature with 200 μl 3% BSA (BioShop Canada Inc. #SKI400.1, lot #9H61850). After washing as above, a four-step, two-fold serial dilution series of patient serum or plasma (0.5–4 μl of sample) was incubated for 1 hr. The wells were washed as above, and incubated with 50 ng biotinylated recombinant ACE2 for 1 hr. After washing as above, the wells were incubated with 44 ng Streptavidin-Peroxidase Polymer (Sigma, #S2438). The resultant signal was developed and quantified with TMB in an identical manner to the direct ELISA assays. Due to day-to-day variation in signal, all OD450 values are normalized to the OD450 of the well where no patient serum/antibody was added for each sample. All values are expressed in this ratio space.

*<u>Variations of this protocol</u>* included using a different source of RBD (BEI Resources NR-52306) and using spike trimer as shown above (670ng/well) (**Figure 1C** and **Supplemental Figure 16**). Another variation of the assay was to bind non-biotinylated ACE2 to the plate (100 ng) and to use biotinylated RBD (50 ng) for detection **(Supplemental Figure 3)**.

### Viral neutralization assays

Neutralization assays on the Canadian Blood Services samples used in **Figure 2** were performed by two independent laboratories, the National Microbiology Laboratory of the Public Health Agency of Canada (NML), and the Wadsworth Center, New York State Department of Health (Wadsworth). The cytopathic effect-reduction neutralization assay on the recombinant GenScript antibody was performed in Toronto.

For the PRNT assay at NML, SARS-CoV-2 (Canada/ON_ON-VIDO-01-2/2020, EPI_ISL_42517) stocks were titrated (7) for use in a plaque reduction neutralization test (PRNT) adapted from a previously described method for SARS-CoV-1 (36). Briefly, serological specimens were diluted 2-fold from 1:20 to 1:640 in DMEM supplemented with 2% FBS and incubated with 50 PFU of SARS-CoV-2 at 37 °C and 5% CO_2_ for 1 hr. The sera-virus mixtures were added to 12-well plates containing Vero E6 cells at 100% confluency, followed by incubation at 37 °C and 5% CO_2_ for 1 hr. After adsorption, a liquid overlay comprised of 1.5% carboxymethylcellulose diluted in MEM, supplemented with 4% FBS, L-glutamine, non-essential amino acids, and sodium bicarbonate, was added to each well and the plates were incubated at 37 °C and 5% CO_2_ for 72 hr. The liquid overlay was removed and the cells were fixed with 10% neutral-buffered formalin for 1 hr at room temperature. The monolayers were stained with 0.5% crystal violet for 10 min and washed with 20% ethanol. Plaques were enumerated and compared to controls. The highest serum dilution resulting in 50% and 90% reduction in plaques compared with controls were defined as the PRNT50 and PRNT90 endpoint titers, respectively. PRNT50 titers ≥1:160 and PRNT90 titers ≥1:20 were considered positive.

For the PRNT assay at Wadsworth, the assay for the detection of SARS-CoV-2 neutralizing antibodies was a modified version of previously described methods (37-39). Patient sera and SARS-CoV-2 (USA/WA-1/2020, BEI Resources, #NR-52281) were diluted in Vero E6 cell culture maintenance medium (Eagle’s Minimal Essential Medium, 2% heat-inactivated fetal bovine serum, 200 U/ml Penicillin G, 200 U/ml Streptomycin). Patient samples were serially diluted 1:10 to 1:320 and mixed with an equal volume of virus containing 150 plaque forming units. Virus and serum mixtures were incubated at 37 °C and 5% CO_2_ for 1 hr. Following the initial incubation, 0.1 mL of each dilution was plated in a single well of a 6 well plate containing confluent monolayers of Vero E6 cells (ATCC, CRL-1586) and allowed to adsorb for 1 hr at 37 °C and 5% CO_2_. Following adsorption, cell cultures were overlaid with 0.6% agar in cell culture medium and returned to the incubator. At two days postinfection, a second overlay containing 0.2% neutral red was added. Monolayers were inspected for two days and plaques were counted. Antibody titers were reported as the inverse of the serum dilution resulting in 50% (PRNT50) and 90% (PRNT90) reduction in plaques as compared to the virus inoculum control.

For the cytopathic effect-reduction neutralization assay in Toronto, 200 μL of 0.2×10^6^ VeroE6 cells/mL were seeded into a 96-well flat bottom plate to adhere overnight. All plasma and serum samples were heat inactivated at 56 °C for 30 min. In a separate 96-well plate, the serum, plasma or antibody (1 μg/ml) samples were serially diluted 2-fold eight times in serum-free DMEM starting from a dilution of 1:20 to 1:2560 in a volume of 25 μL. To all wells, 25 μL of SARS-CoV-2 SB2 Clone 1 was added ensuring that each well had a dose of 100 TCID. For the cell control, 50 μL of serum free DMEM was added. For the virus control, 25 μL of SARS-CoV-2 SB2 Clone 1 was added with a dose of 100 TCID, and topped off with 25 μL of serum free DMEM. The plate was incubated for 1 hr at 37 °C, 5% CO_2_ with shaking every 15 min. After incubation, all the media from the VeroE6 culture was removed, and the full 50 μL of serum/SARS-CoV-2 co-culture was layered on the cells. The plate was again incubated for 1 hr at 37 °C, 5 % CO_2_, with shaking every 15 min. After the incubation, the inoculum was removed and 200 μL of DMEM containing 2 % FBS was added. The plate was incubated for 5 days and cytopathic effect (CPE) was tracked.

### Lentiviral spike pseudotyping assay

The assay was established using constructs previously described (8) (constructs obtained through a kind gift from Jesse Bloom and Katharine Crawford, Fred Hutchison Cancer Research Centre, Seattle, WA, USA, and now available through BEI Resources), and optimized in-house. Major changes to the reported protocol included: 1) Use of a 2^nd^ generation psPAX2 (Addgene, #12260) lentivirus packaging system instead of the 3^rd^ generation system used by the Bloom lab, 2) Production of spike pseudotyped virus-like particles (VLPs) at 33°C, 3) A neutralization assay plate layout that increases throughput, 4) Adjustments to the luciferase protocol to minimize variability in readings and, 5) Use of a cell line that co-expresses ACE2 and TMPRSS2. To generate this cell line, entry vectors for ACE2 and TMPRSS2 coding sequences were cloned into pLenti CMV Puro DEST (Addgene, #17452) and pLenti CMV Hygro DEST (Addgene, #17454) respectively. The resulting transfer vectors were used to generate lentivirus via the 2^nd^ generation psPAX2 and VSV-G (Addgene, #8454). HEK293T cells were transduced with ACE2 lentivirus at an MOI <1 and selected with puromycin (1 μg/mL) to generate a stable population. These cells were subsequently transduced with TMPRSS2 lentivirus and selected with hygromycin (200 μg/mL) in a similar fashion.

For VLP generation, HEK293T cells were transiently co-transfected in a 6-well plate format containing 2 ml growth medium (10% FBS, 1% Pen/Strep in DMEM) with 1.3 μg psPAX2, 1.3 μg pHAGE-CMV-Luc2-IRES-ZsGreen-W (BEI, #NR-52516; a kind gift from Jesse Bloom and Katharine Crawford; lentiviral backbone plasmid that uses a CMV promoter to express luciferase followed by an IRES and ZsGreen) and 0.4 μg HDM-IDTSpike-fixK (BEI, #NR-52514; a kind gift from Jesse Bloom and Katharine Crawford; expresses under a CMV promoter a codon-optimized Wuhan-Hu-1 spike (Genbank, NC_045512) using 8 μl JetPrime in 500 μl JetPrime buffer. After 8 hr of transfection, the medium was replaced by 3 mL of DMEM containing 5% heat-inactivated FBS, 1% Pen/Strep and the cells incubated for 16 hr at 37 °C and 5% CO_2_ and then transferred to 33 °C and 5% CO_2_ for an additional 24 hr. At 48 hr post transfection, the supernatant was collected, spun at 500*g* for 5 min, filtered through a 0.45 μm filter and frozen at −80 °C. The virus titers were evaluated using HEK293T-ACE2/TMPRSS2 cells at 10K cells per well on a Poly-L-Lysine [5-10 μg/mL] coated 96-well plate using HI10 media (10% heat-inactivated FBS, 1% Pen/Strep) and a virus dilution resulting in >1000 relative luciferase units (RLU) over control (~1:100 virus stock dilution).

For the neutralization assay, 2.5-fold serial dilutions of the serum samples were incubated with diluted virus at a 1:1 ratio for 1 hr at 37 °C before being transferred to plated HEK293-ACE2/TMPRSS2 cells and incubated for an additional 48 hr at 37 °C and 5% CO_2_. After 48 hr, cells were lysed and Bright-Glo luciferase reagent (Promega, #E2620) was added for 2 min before reading with a Perkin-Elmer Envision instrument.

### Data analysis and figure generation

Area Under the Curve (AUC) values were tabulated for both the direct binding ELISA and the snELISA using R version 4.0.1 and R package pracma. For the snELISA, the ratios (normalized values) are used in the AUC calculations. To identify outliers, we calculated the distance of each point from the regression line using total least squares and labelled points with distances > 0.4.

For the lentiviral pseudotyping assays, 50% inhibitory concentration or dilution (IC50 or ID50) were calculated with non-linear regression [log(inhibitor) vs. normalized response - Variable slope] using GraphPad Prism 8 (GraphPad Software Inc., San Diego, CA, USA).

For the extended direct binding dilution series, titres were calculated by taking the dilution of serum that produced 50% of the maximum response in the ELISA as determined by the non-linear regression line [Sigmoidal, 4PL, X is log(dilution)] using GraphPad Prism 8.

The assay reproducibility was estimated across experiments by comparing the AUC values for those samples profiled across different batches.

CBS13 (n=3) CV for displacement was 5.1% and direct binding was 5.5%; CBS16 (n=3) CV for displacement was 3.4% and 11.5% for binding;

CBS50 (n=2) CV for displacement was 9.9% and binding was 0.7%

### Statistics

When applicable, graphical data from experiments with three or more replicates are presented as mean ± SEM.

### Study approval

All samples were collected after Research Ethics Board (REB) review. The ELISA assays were performed at the Lunenfeld-Tanenbaum Research Institute with Mount Sinai Hospital (MSH; Toronto, ON) Research Ethics Board (REB) approval (study number: 20-0078-E). External samples were transferred through Material Transfer Agreements. All research has been performed in accordance with relevant guidelines and regulations. All participants have provided informed consent. The samples were de-identified prior to transfer to the assay laboratory.

## Author contributions

KTA and ACG designed the snELISA and the direct RBD antibody assay

ZL and JMR designed the protein expression, biotinylation and purification procedures

RS and PST optimized the lentiviral pseudotyping assay

EJM, HW, MAD, AD, RG, KAM, PB and MO developed, performed and analyzed the PRNT and/or cytopathic effect-reduction neutralization assays KTA and BR performed direct ELISA experiments

YD designed the VHH72hFc1X7 construct, expression and purification procedures

JHW and MBR implemented the automated direct binding ELISA

KC helped coordinate the project

SD provided samples, coordinated neutralization testing and integrated PRNT and snELISA data KTA and ACG analyzed the snELISA data

JLG, AJM, SM, MO and SD contributed essential patient samples KTA, JMR and ACG wrote the manuscript with input from all authors The order of authors for the two co-first author was determined by the contribution of KTA in the overall study design, and data analysis and manuscript preparation.

## Acknowledgements

We thank Janet McManus at Canadian Blood Services for her technical and logistical expertise and the Wadsworth Center Media and Tissue Culture Core. We thank Joan Wither for the lupus patient samples, and Jesse Bloom and Katharine Crawford for sharing protocols and reagents for the lentiviral S pseudotyping assay. The following reagent was produced by Florian Krammer’s group under HHSN272201400008C and obtained through BEI Resources, NIAID, NIH: Spike Glycoprotein Receptor Binding Domain (RBD) from SARS-Related Coronavirus 2, Wuhan-Hu-1, Recombinant from HEK293 Cells, NR-52306. We thank Matthew Stuible and Alex Pelletier (National Research Council; NRC) for VHH72hFc1X7 expression and purification, Joe Schrag (NRC) for SEC-UPLC/MALS analysis and members of our laboratories and the Network Biology Collaborative Centre for advice and technical help. Funding for the development of the assays in the Gingras lab was provided through generous donations from the Royal Bank of Canada (RBC), QuestCap and the Krembil Foundation to the Sinai Health System Foundation. We also acknowledge an “Ontario Together” team grant from the Government of Ontario to JG, JMR, ACG, MO, AM to develop serology and saliva assays. The equipment used is housed in the Network Biology Collaborative Centre at the Lunenfeld-Tanenbaum Research Institute, a facility supported by Canada Foundation for Innovation funding, by the Ontarian Government and by Genome Canada and Ontario Genomics (OGI-139). We also acknowledge funding support from New York State Department of Health (to KAM). AJM, JMR and ACG are supported by the Canadian Institutes of Health Research (CIHR 439999, FRN 162305 and FDN 143301, respectively). ACG is also supported by a Canada Research Chair, Tier 1, in Functional Proteomics. KTA is a recipient of an Ontario Graduate Scholarship.

## Supplemental Figures and legends

**Supplemental Figure 1.**
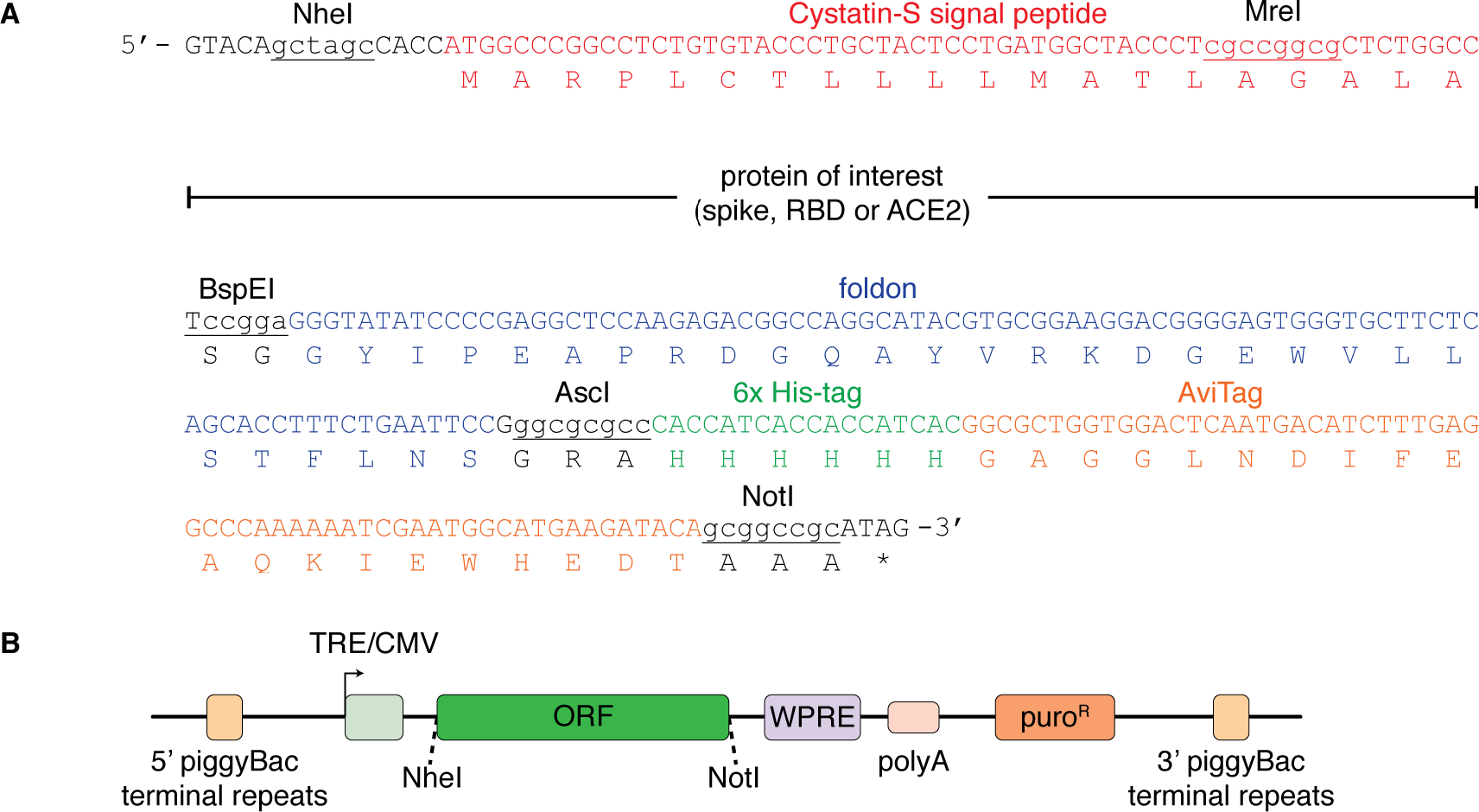
Details of the protein expression constructs. (**A**) Multiple cloning site and fusion tags. (**B**) Schematic of the expression vectors used to express recombinant ACE2, spike and RBD.

**Supplemental Figure 2.**
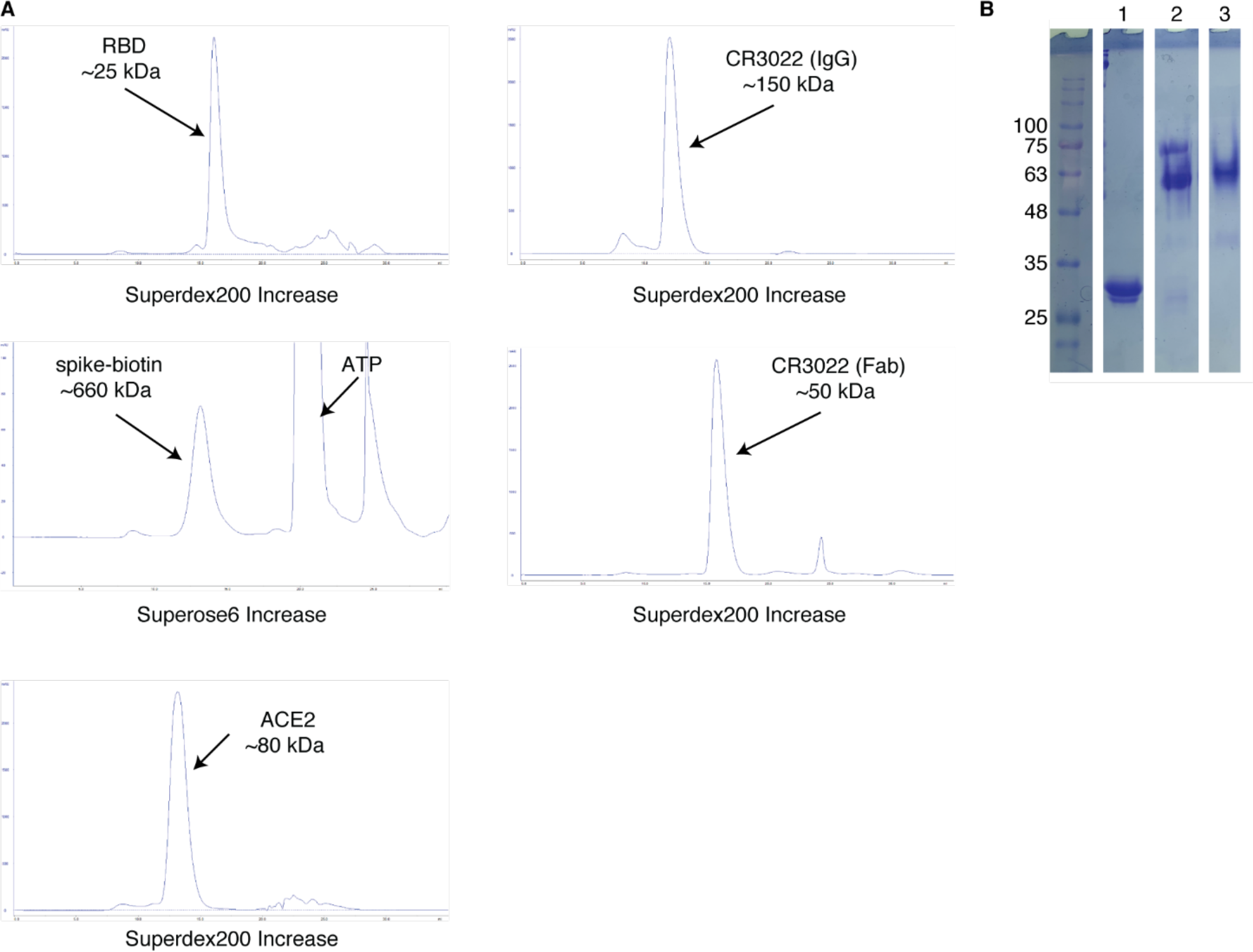
Protein purification and biotinylation. (**A**) Size-exclusion chromatography profiles of the RBD, ACE2 and the CR3022 Fab/IgG run on a Superdex 200 Increase column. The spike ectodomain trimer was run on a Superose 6 Increase column. Purification of the spike after the biotinylation reaction is shown to illustrate the separation of the protein from the small molecules. (**B**) Band-shift assay to assess RBD biotinylation: lane 1, SARS-CoV-2 RBD-biotin; lane 2, SARS-CoV-2 RBD-biotin + streptavidin; lane 3, streptavidin.

**Supplemental Figure 3.**
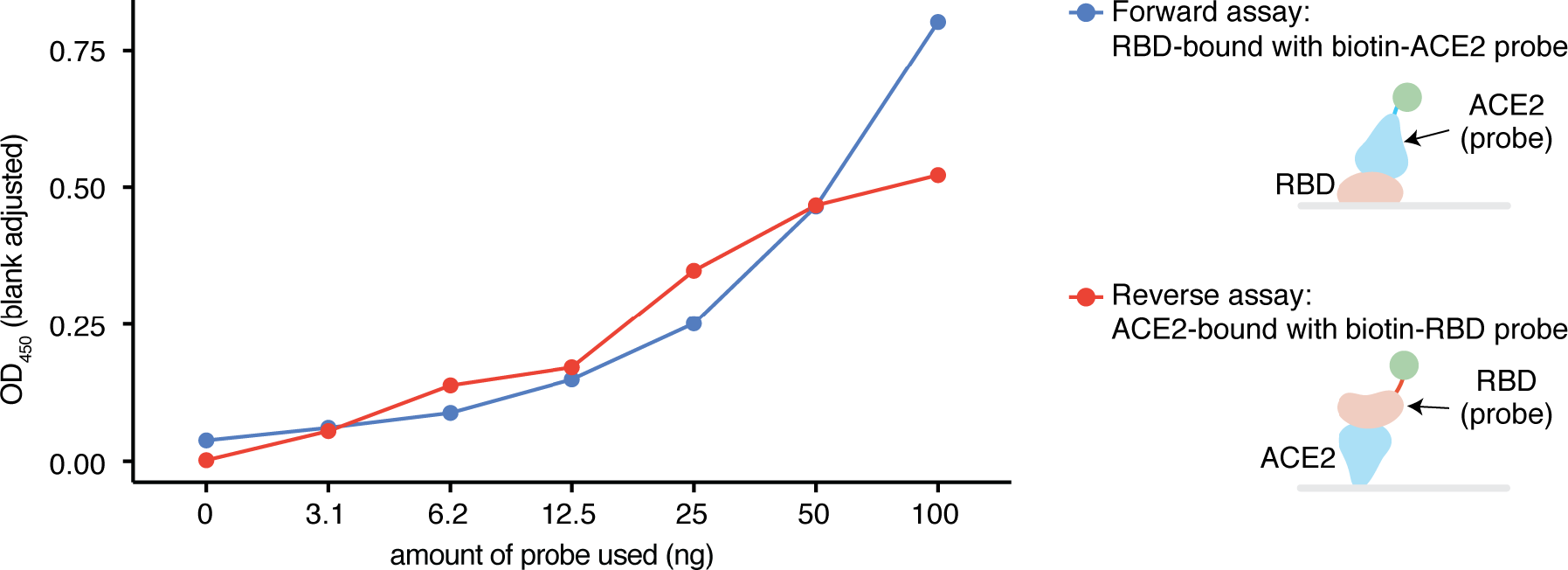
Comparison of the signal of the RBD-bound (forward) and ACE-bound (reverse) assays. The forward assay (blue) is where non-labelled RBD is adsorbed onto the ELISA plate and biotinylated ACE2 is added as a probe. The reverse assay (red) is where non-biotinylated ACE2 is adsorbed and probed with biotinylated RBD. After the addition of the probing protein, the presence of the biotinylated protein is evaluated with poly-HRP and TMB. This seven-point curve was generated from one experiment.

**Supplemental Figure 4.**
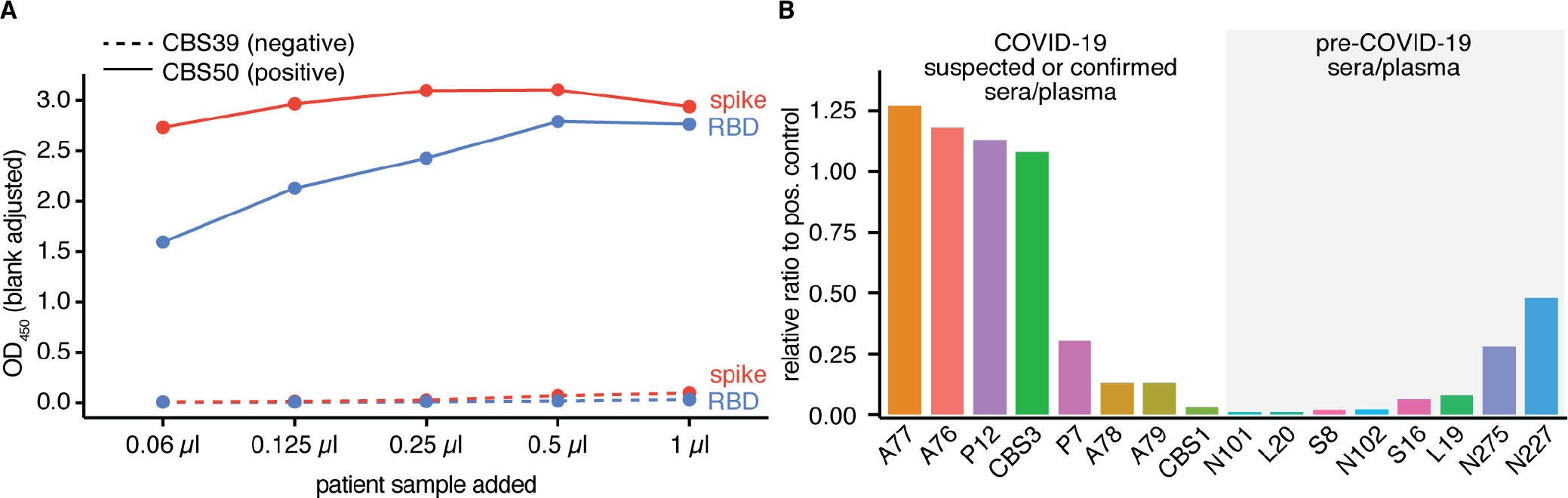
Direct binding ELISAs for assay optimization. (**A**) Direct binding ELISA probing for IgG on a dilution series of serum samples CBS39 (negative) and CBS50 (positive). Related to **Figure 1E**. This five-point curve was generated from one experiment. (**B**) Single-point direct binding ELISA assay of 12 pilot samples on the RBD. Data expressed as a ratio to the OD_450nm_ of a convalescent, positive plasma sample used for normalization (P1). The bars represent the mean value from two technical replicates from one experiment. Related to **Figure 1F**.

**Supplemental Figure 5.**
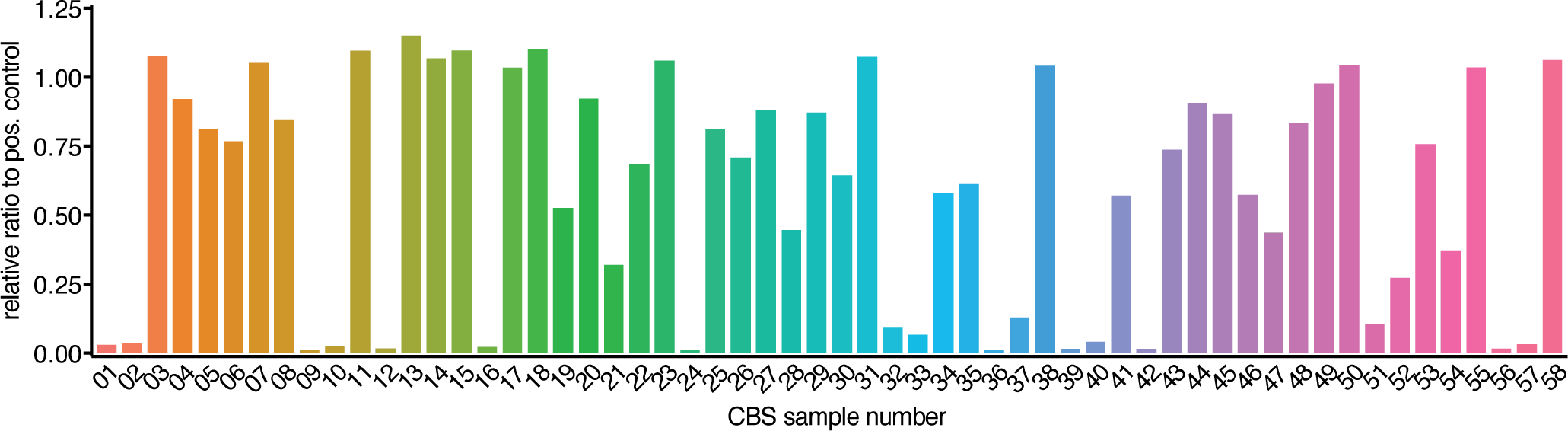
Single-point direct binding ELISA assay of all CBS samples on the RBD. Data expressed as a ratio to the OD_450nm_ of a convalescent, positive plasma sample used for normalization (P1). The bars represent the mean from two technical replicates from one experiment (n = 58).

**Supplemental Figure 6.**
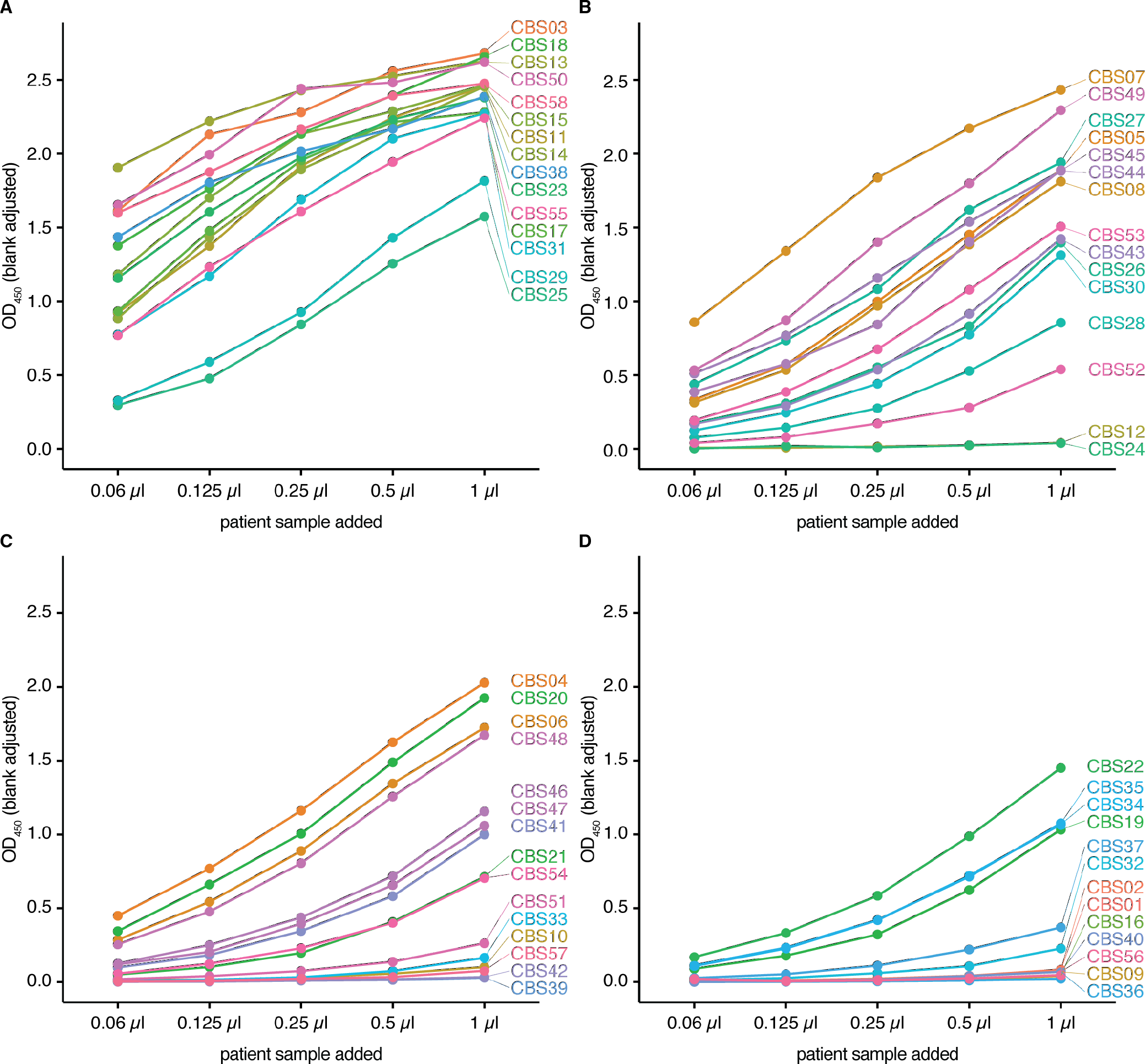
The entire set of direct binding ELISAs with titrations on different samples of the Canadian Blood Services cohort. Samples were ranked based on the AUC of the snELISA curve and were plotted in four batches starting with samples most effective at blocking the RBD-ACE2 interaction (**A**), to the lowest (**D**). Related to **Figure 2B** and **Supplemental Figure 8**. The five-point curves were generated from one experiment (n = 58).

**Supplemental Figure 7.**
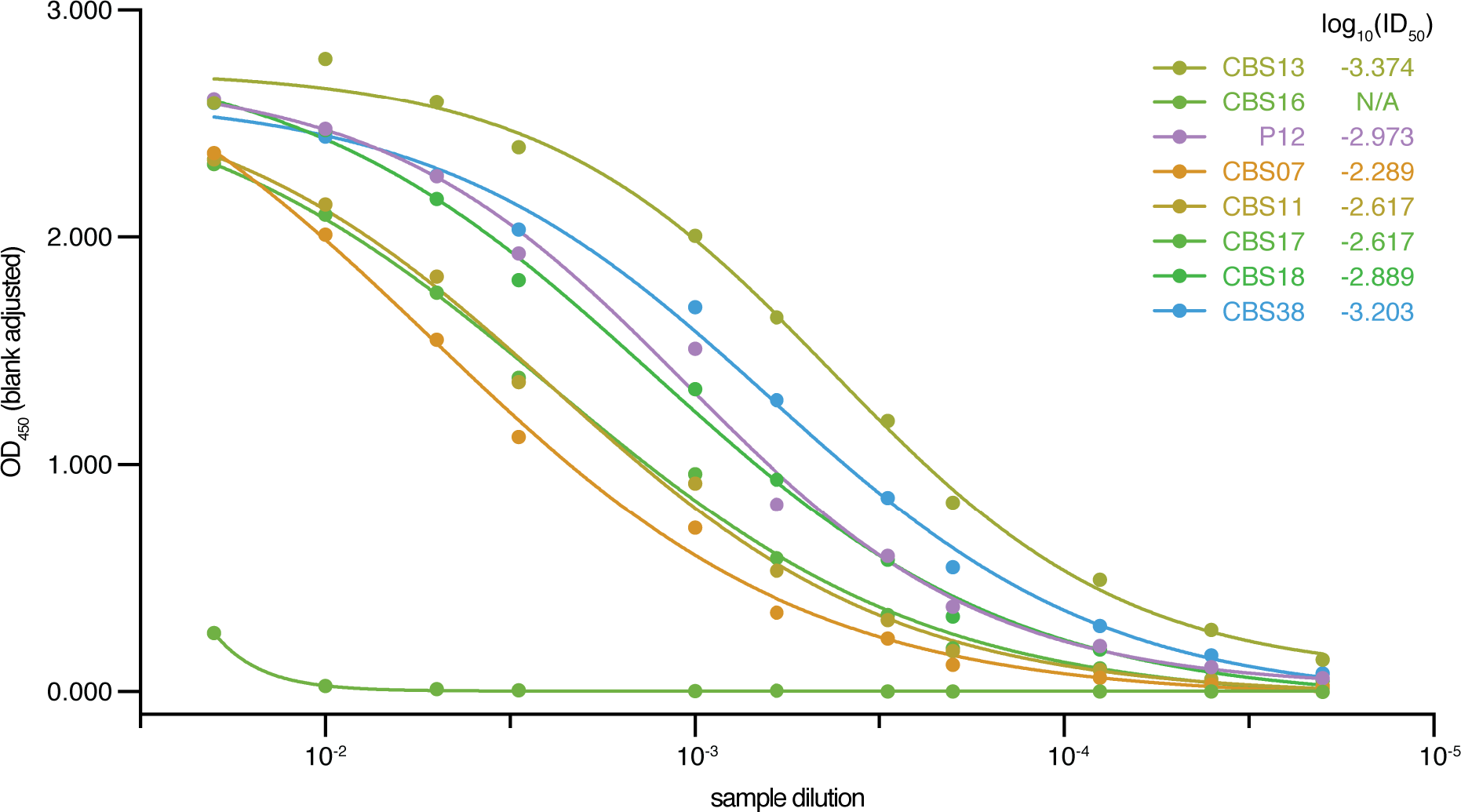
An extended direct binding assay dilution series. Six CBS samples exhibiting a high OD_450nm_ at the lower end of the titration curve in the direct binding assay (0.06 μl) were further diluted to generate an extended direct binding titration curve. Related to **Figure 2A** and **Supplemental Figure 6**. The 11-point dilution curves were generated from one experiment.

**Supplemental Figure 8.**
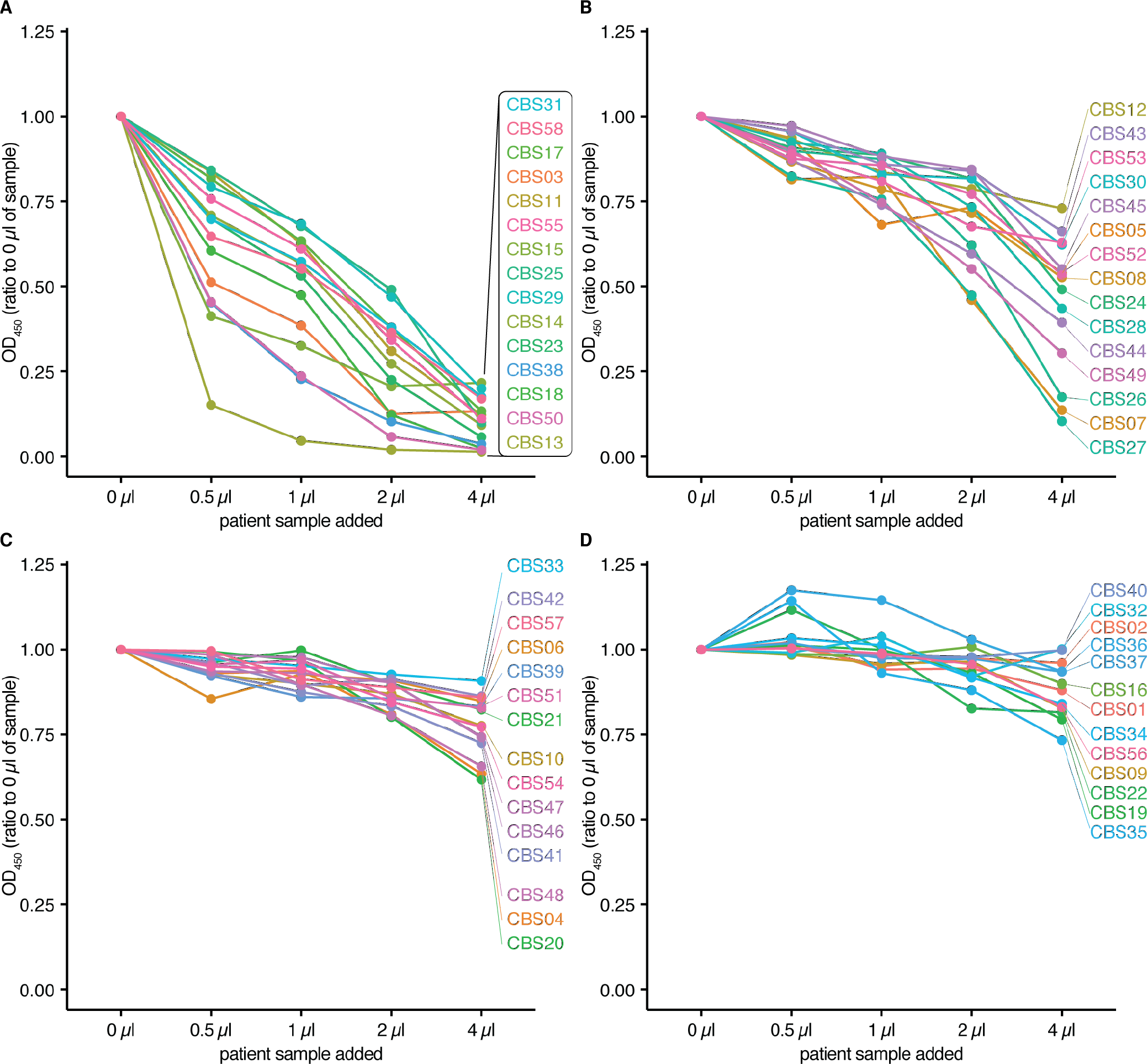
The entire set of snELISAs with titrations on different samples of the Canadian Blood Services cohort. Samples were ranked based on the AUC of the snELISA curve and were plotted in four batches starting with samples most effective at blocking the RBD-ACE2 interaction (**A**), to the lowest (**D**). Related to **Figure 2B**. The 5-point dilution curves were generated from one experiment (n = 58).

**Supplemental Figure 9.**
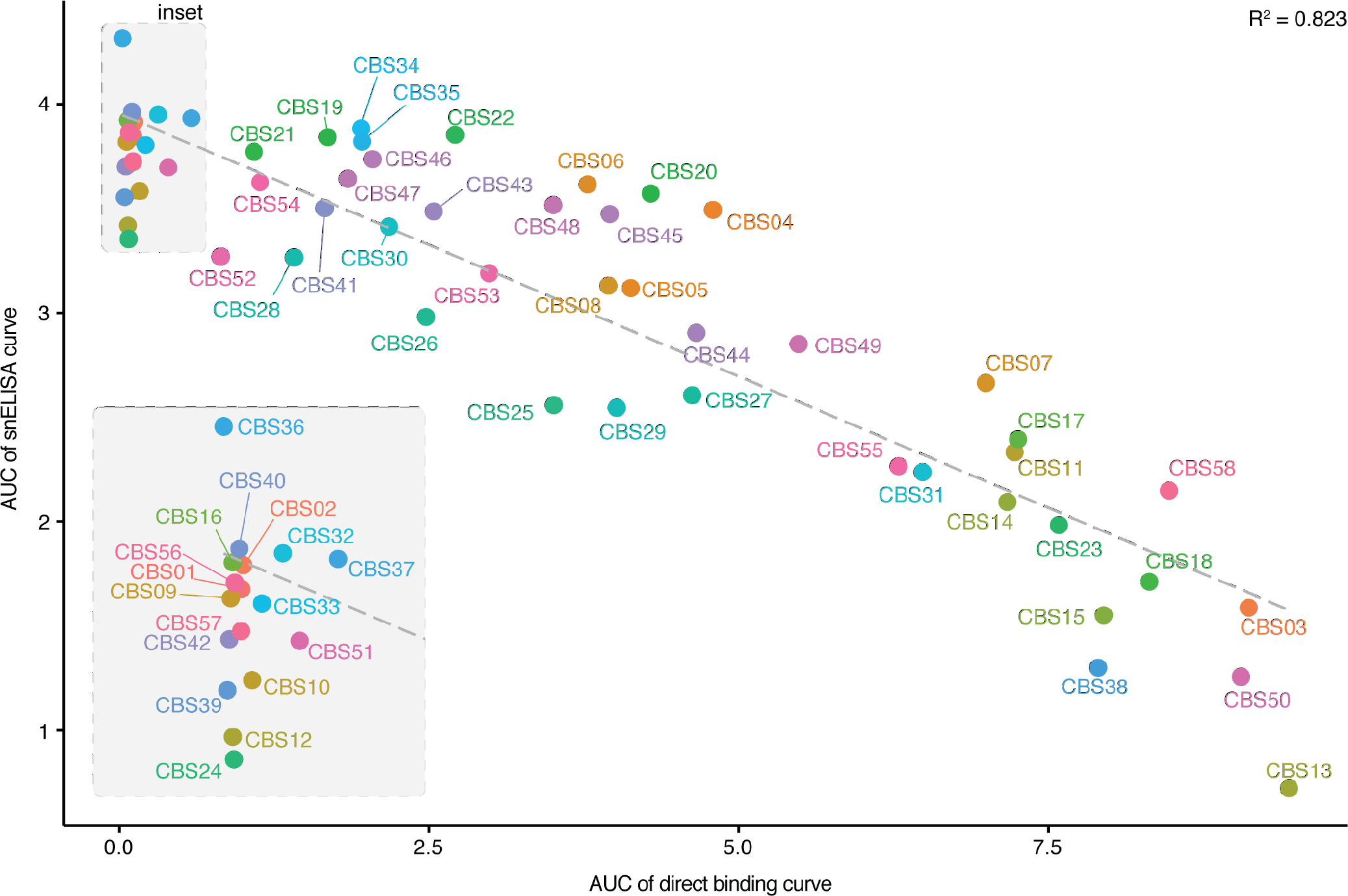
Complete correlation plot between the AUCs for the direct and surrogate neutralization ELISAs for all samples profiled with all of the samples labelled. Related to **Figure 2C**.

**Supplemental Figure 10.**
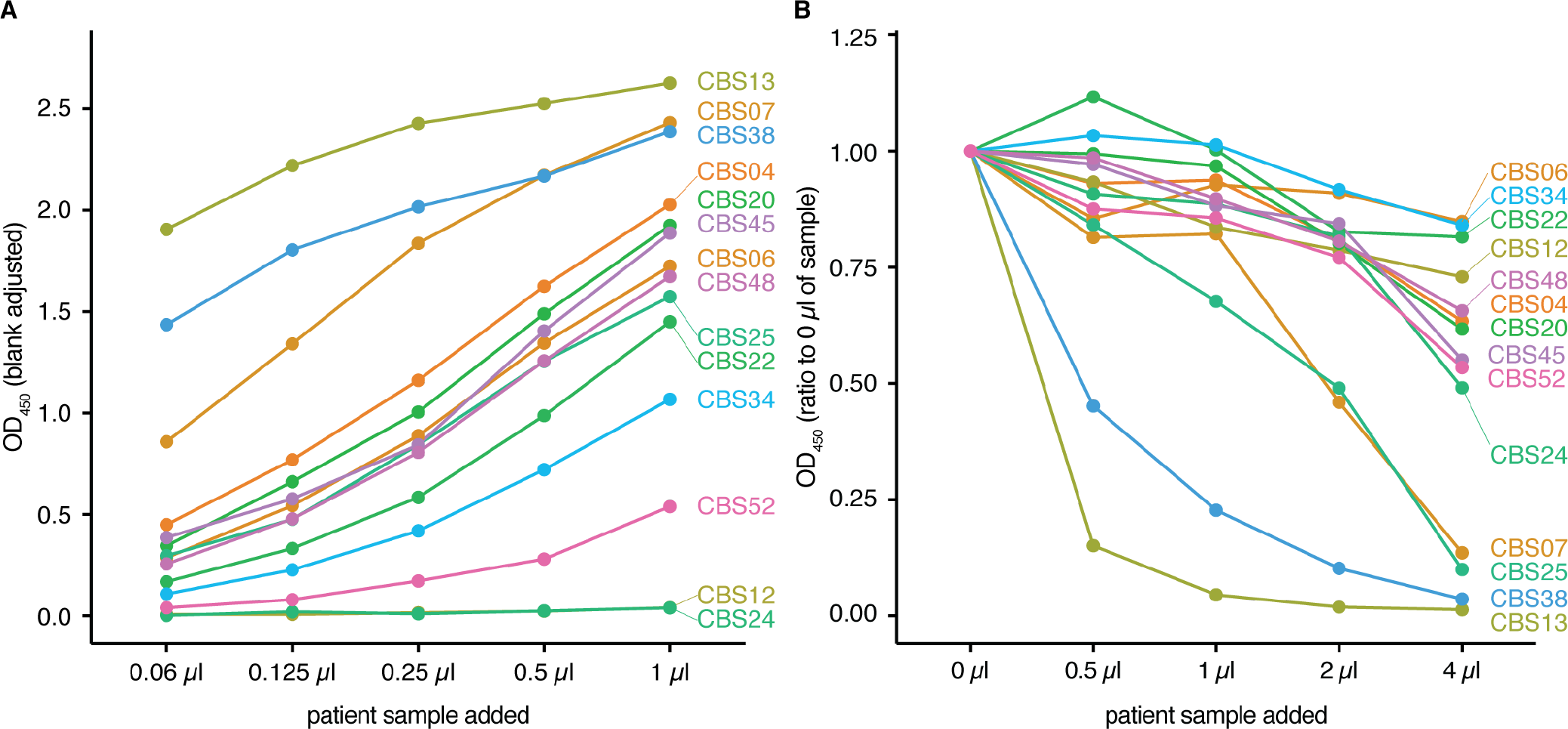
ELISA results of CBS samples labelled as outliers (TLS error >0.4). (**A**) Direct binding assay titration curves and (**B**) snELISA titration curves.

**Supplemental Figure 11.**
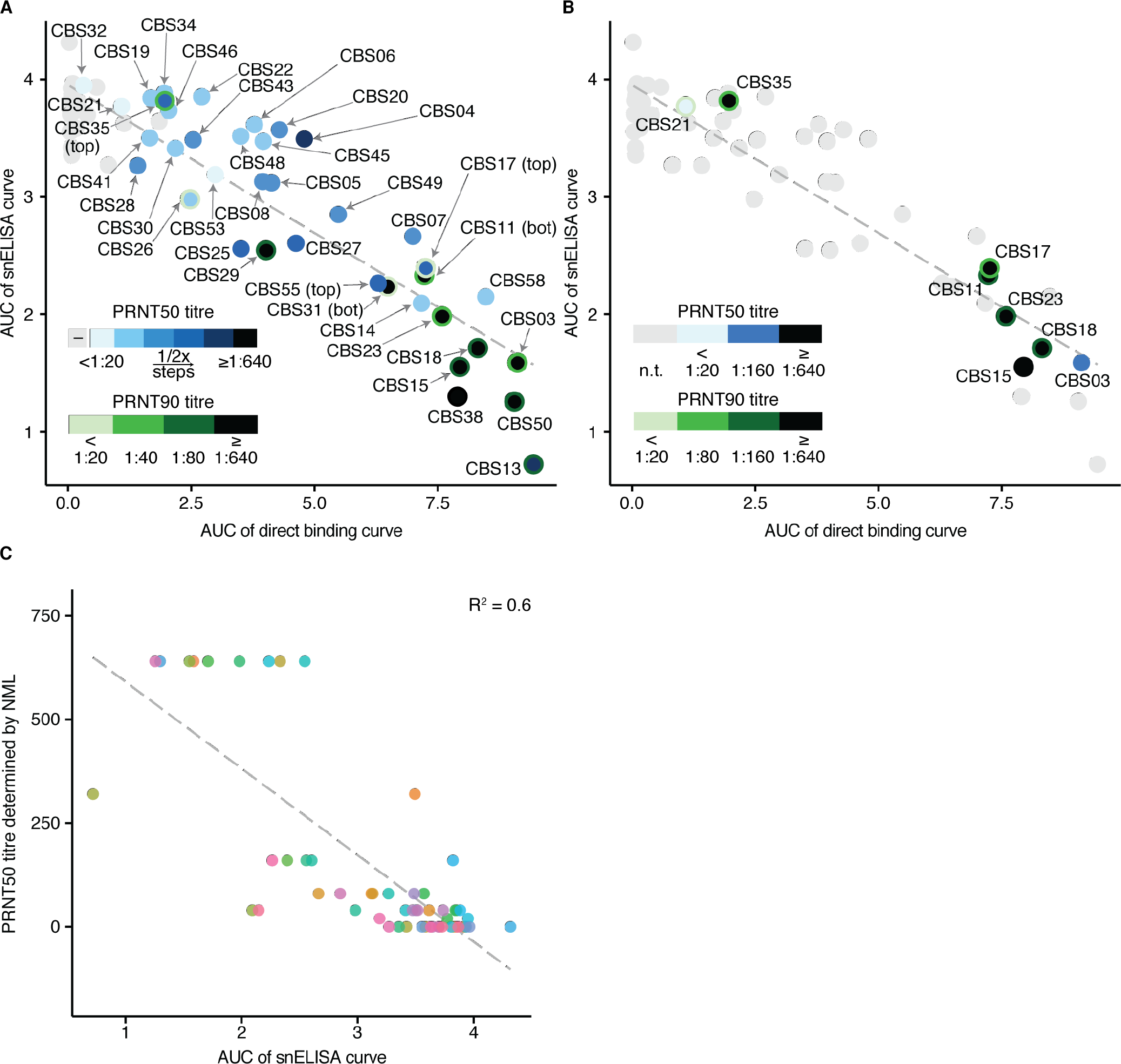
PRNT50 and PRNT90 data overlaid on the AUC correlation plot. (**A**) The correlation curve from **Figure 2C** with PRNT50 and PRNT90 data from the NML overlaid on each data point (n = 57). The point fill (blue) represents the PRNT50 titre whereas the outline stroke colour (green) represents the PRNT90 titre. Grey points indicate samples that were negative with both PRNT50 and PRNT90. (**B**) Same as panel (**A**) but with data from Wadsworth (n = 8; samples in grey were not tested). (**C**) Correlation between the results of the PRNT50 titres determined by the NML and the AUC of the snELISA curves calculated for the CBS samples. Note that the titres are binned in 2-fold increments starting with <1:20 and ending with ≥1:640. The grey line represents the linear regression for n = 57 samples. This figure is related to **Figure 2C**.

**Supplemental Figure 12.**
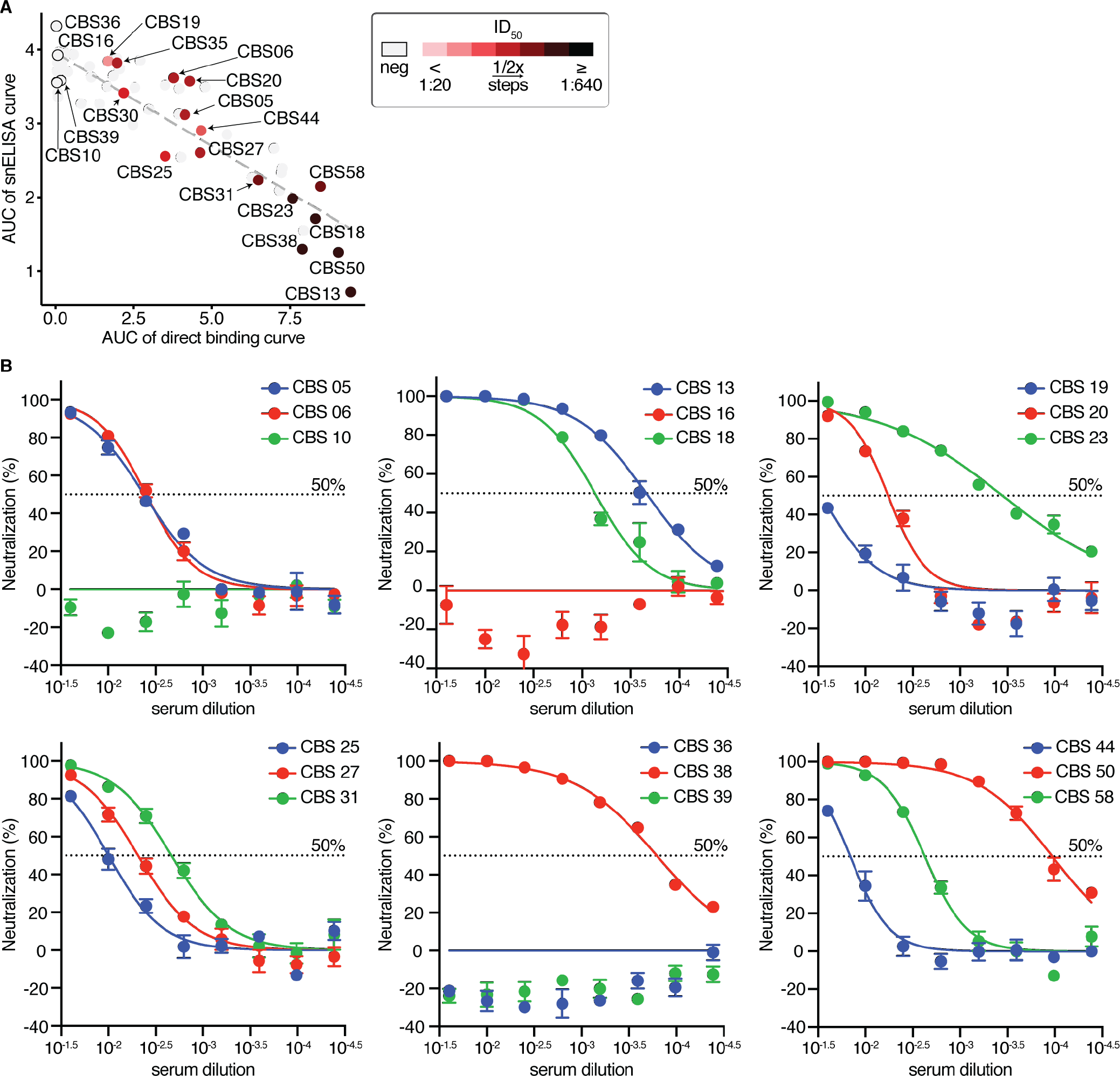
Lentiviral spike pseudotyped assay on selected CBS samples. (**A**) The indicated samples were tested by the lentiviral spike pseudotyped assay (n = 21). The ID50 of tested samples were calculated and overlaid on the snELISA. Related to **Figure 2C**. (**B**) Neutralization curves used to calculate the ID50 values of each sample in **A** (n = 18). Error bars represent the standard error of the mean (SEM) of three replicates.

**Supplemental Figure 13.**
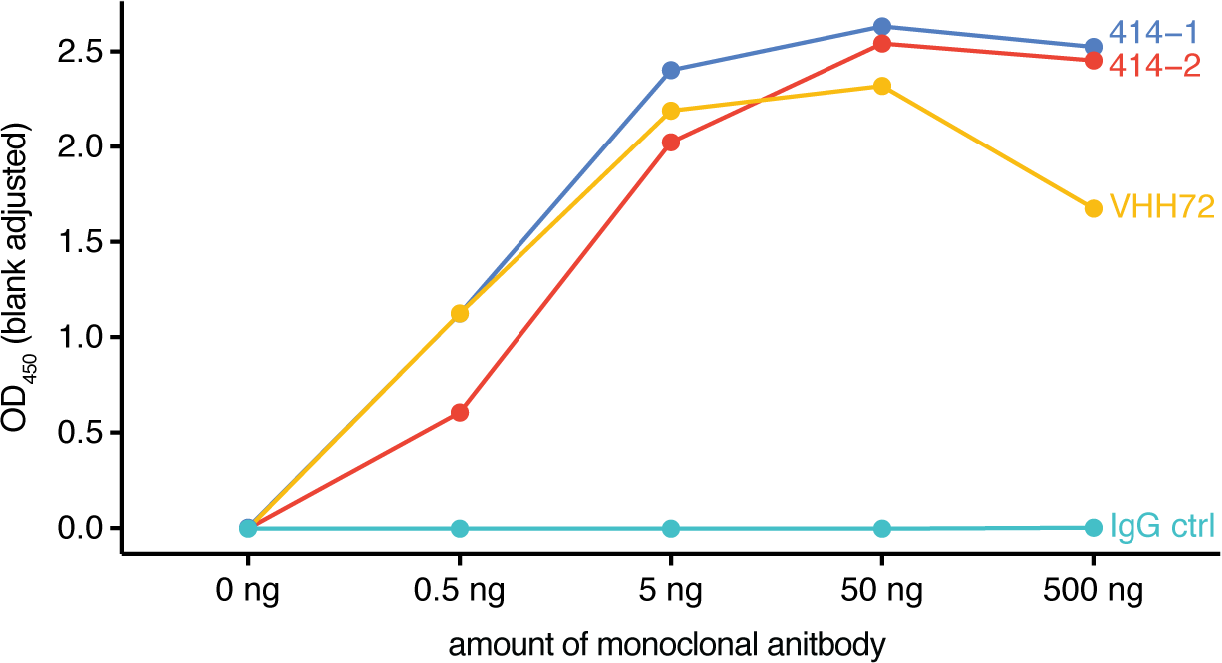
Direct binding assay on monoclonal antibodies. Direct binding ELISA probing for IgG on a dilution series of commercial monoclonal antibodies. The 5-point dilution curves were generated from one experiment. Related to **Figure 3C**.

**Supplemental Figure 14.**
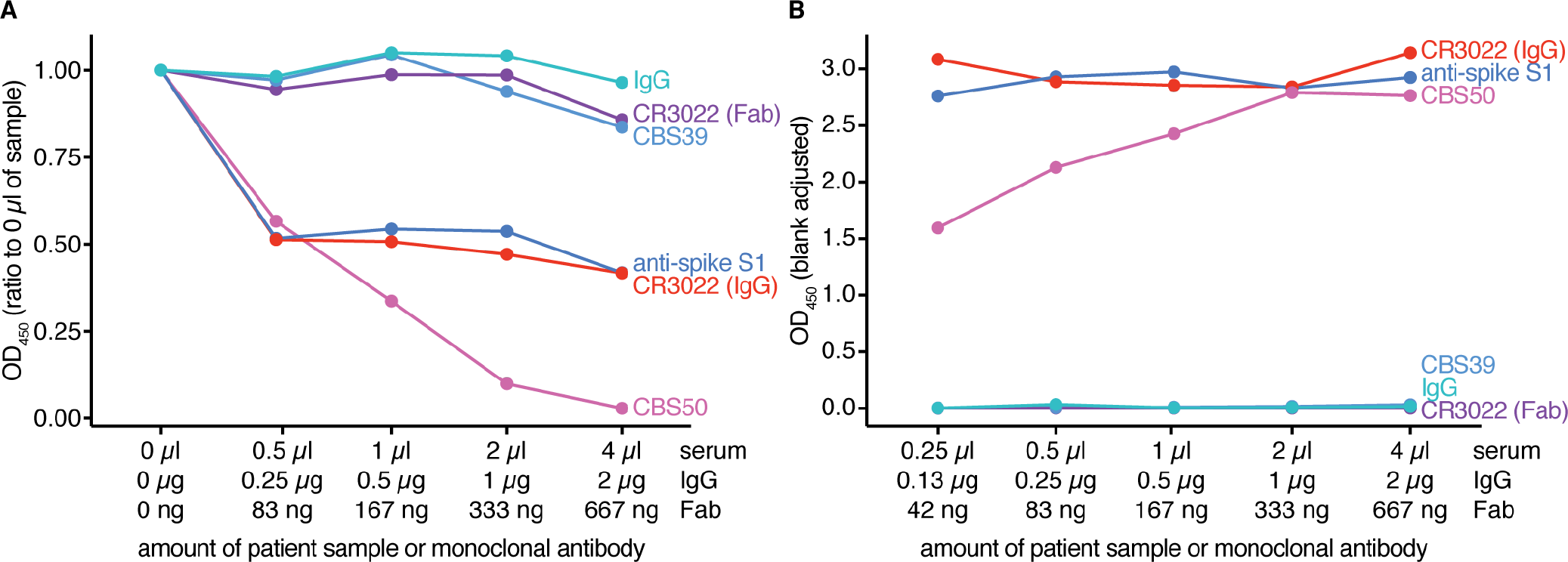
Surrogate neutralization and direct binding ELISAs of monoclonal antibodies and affinity reagents. (**A**) snELISA of patient samples and monoclonal antibodies. (**B**) Direct binding assay of a dilution series of the serum samples and antibodies used in **A**. Note that the secondary antibody is an anti-Fc IgG, so the CR3022 (Fab, purple) is not expected to show activity. The 5-point dilution curves were generated from one experiment.

**Supplemental Figure 15.**
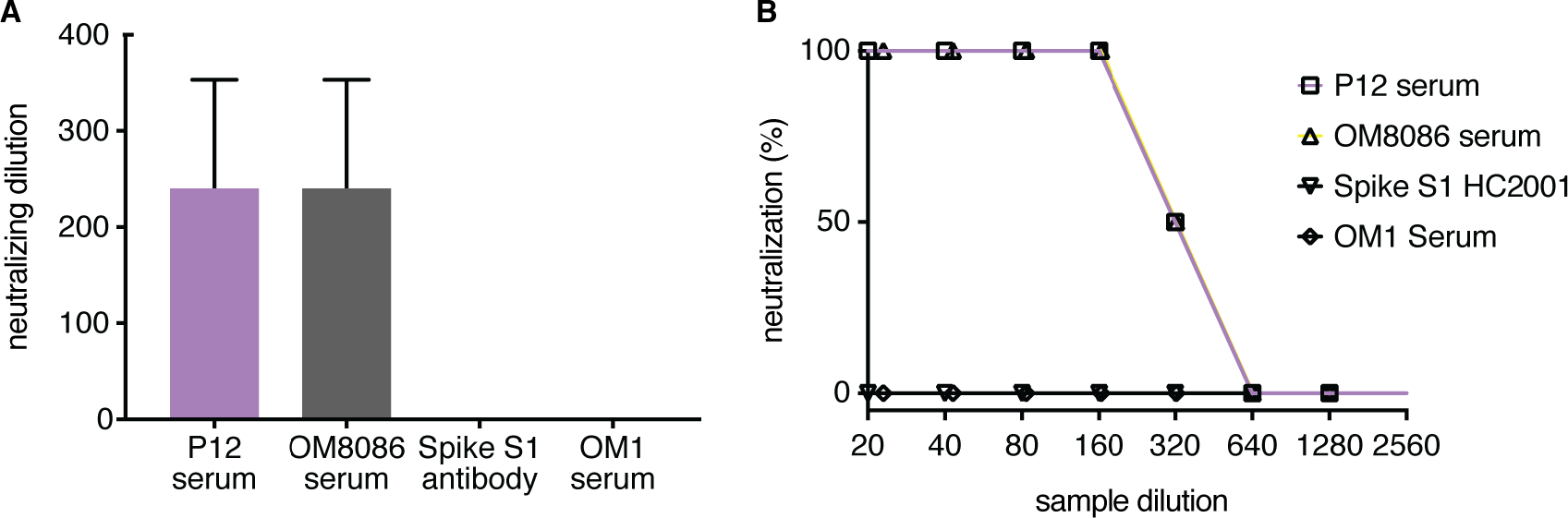
Cytopathic effect-reduction neutralization assay on the GenScript antispike S1 antibody. Convalescent sera P12 and OM8086 as well as SARS-CoV-2 unexposed serum OM1 and anti-spike monoclonal antibody HC2001 were serially diluted and co-cultured with SARS-CoV-2. The co-cultures were layered onto VeroE6 cells for 1 hr and plaques were monitored. Samples were run in quadruplicates and error bars represent the SEM. (**A**) neutralization dilution; (**B**) percentage neutralization at indicated dilution.

**Supplemental Figure 16.**
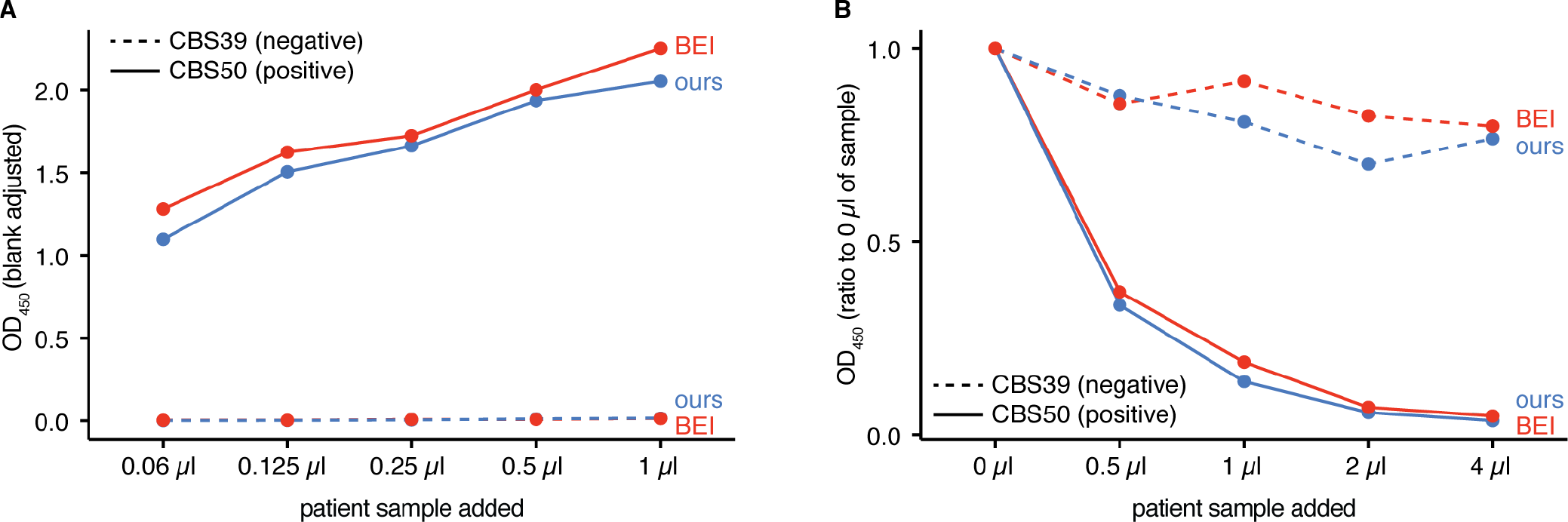
Compatibility of the assay with the RBD from a different source (F. Krammer via BEI). Direct binding ELISA probing for IgG on a dilution series of serum samples CBS39 (negative reactivity) and CBS50 (positive reactivity). The 5-point dilution curves were generated from one experiment, and samples were run in technical duplicates.

**Supplemental Figure 17.**
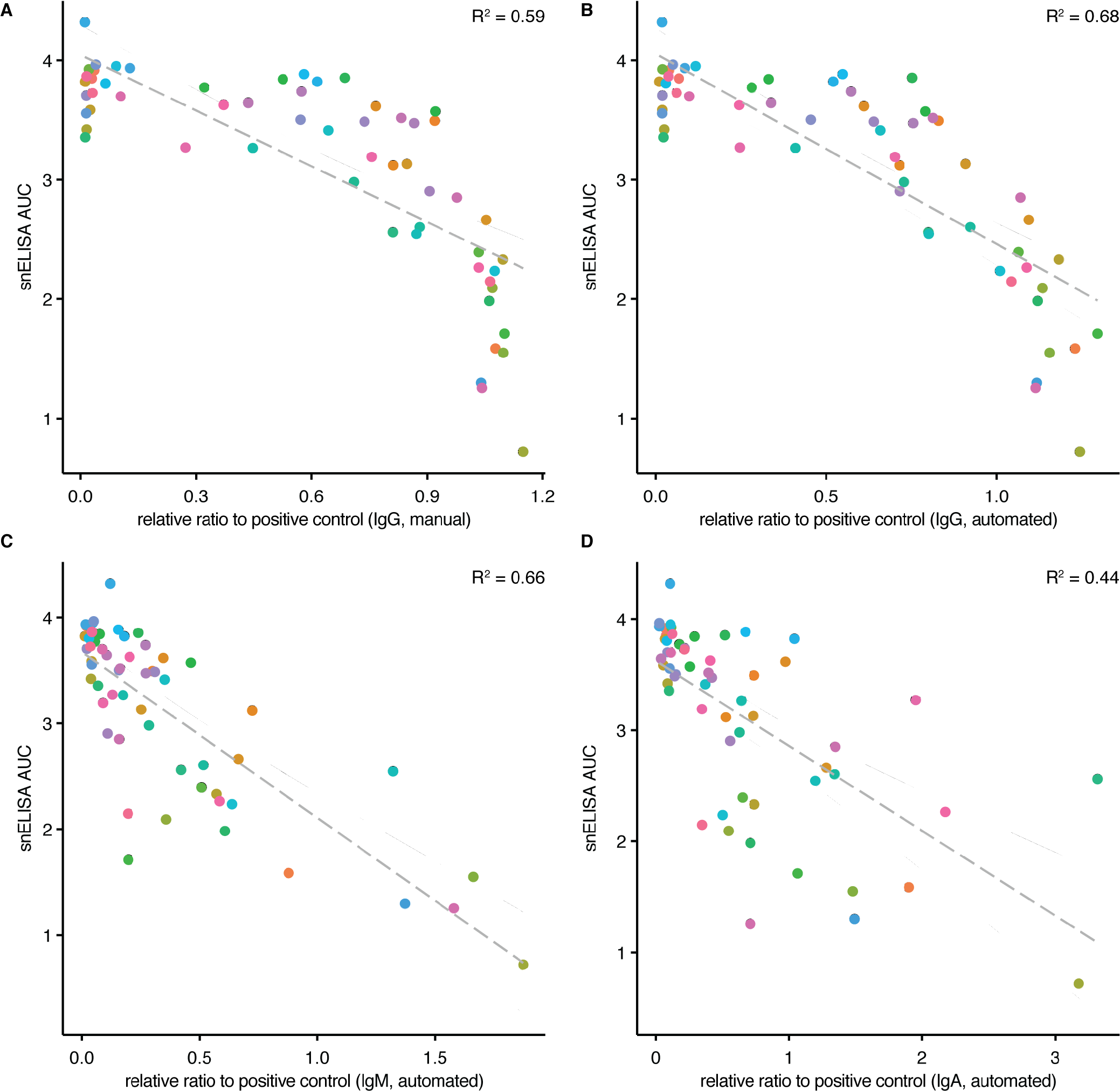
Correlation between the results of the snELISA and direct binding ELISAs. The snELISA AUC values were plotted against the single-point direct binding ELISA results probing for (**A**) IgG using data from manual experiments. Panels (**B-D**) show correlations between snELISA AUCs and single-point direct binding results when samples were probed for IgG, IgM and IgA, respectively, using data from ELISAs conducted with an automated platform and a chemiluminescent readout as in (11). Samples (n = 58) were run in technical duplicates, and the grey line represents the linear regression.

